# Mild neonatal hypoxia disrupts adult hippocampal learning and memory and is associated with CK2-mediated dysregulation of synaptic calcium-activated potassium channel KCNN2

**DOI:** 10.1101/2024.07.10.602558

**Authors:** Art Riddle, Taasin Srivastava, Kang Wang, Eduardo Tellez, Hanna O’Neill, Xi Gong, Abigail O’Niel, Jaden A Bell, Jacob Raber, Matthew Lattal, James Maylie, Stephen A. Back

**Affiliations:** Oregon Health & Science University, Department of Pediatrics; Behavioral Neuroscience; Obstetrics and Gynecology

## Abstract

**Objective:** Although nearly half of preterm survivors display persistent neurobehavioral dysfunction including memory impairment without overt gray matter injury, the underlying mechanisms of neuronal or glial dysfunction, and their relationship to commonly observed cerebral white matter injury are unclear. We developed a mouse model to test the hypothesis that mild hypoxia during preterm equivalence is sufficient to persistently disrupt hippocampal neuronal maturation related to adult cellular mechanisms of learning and memory.

Methods: Neonatal (P2) mice were exposed to mild hypoxia (8%O_2_) for 30 min and evaluated for acute injury responses or survived until adulthood for assessment of learning and memory and hippocampal neurodevelopment.

**Results:** Neonatal mild hypoxia resulted in clinically relevant oxygen desaturation and tachycardia without bradycardia and was not accompanied by cerebral gray or white matter injury. Neonatal hypoxia exposure was sufficient to cause hippocampal learning and memory deficits and abnormal maturation of CA1 neurons that persisted into adulthood. This was accompanied by reduced hippocampal CA3-CA1 synaptic strength and LTP and reduced synaptic activity of calcium-sensitive SK2 channels, key regulators of spike timing dependent neuroplasticity, including LTP. Structural illumination microscopy revealed reduced synaptic density, but intact SK2 localization at the synapse. Persistent loss of SK2 activity was mediated by altered casein kinase 2 (CK2) signaling.

**Interpretation:** Clinically relevant mild hypoxic exposure in the neonatal mouse is sufficient to produce morphometric and functional disturbances in hippocampal neuronal maturation independently of white matter injury. Additionally, we describe a novel persistent mechanism of potassium channel dysregulation after neonatal hypoxia. Collectively our findings suggest an unexplored explanation for the broad spectrum of neurobehavioral, cognitive and learning disabilities that paradoxically persist into adulthood without overt gray matter injury after preterm birth.

## Introduction

Preterm birth is associated with significant neurological complications related to periventricular hypoxic-ischemic white matter injury, which results in the motor disabilities of cerebral palsy.^1,2^ Although advances in neonatal intensive care have led to a decline in the incidence of cerebral palsy to ∼5-10%,^3–5^ the overall disability rate in children has remained stable, because of a paradoxical persistence of cognitive and learning disabilities in up to 50% of preterm survivors. Notably, disturbances in episodic and working memory^6–8^ persist into adulthood^4,5,9^ and are associated with unexplained disturbances in hippocampal growth and development. Most of these children lack MRI-defined hippocampal lesions typical of hypoxic-ischemic injury, which suggests that an alternative mechanism unrelated to hypoxia-ischemia may contribute to memory impairment. Although numerous experimental studies have demonstrated that isolated exposure to hypoxia does not cause neonatal brain injury, recurrent severe neonatal hypoxia was found to cause hippocampal injury.^10,11^ Given these conflicting observations, we assessed whether milder exposure to hypoxia may result in memory impairment related to maturational disturbances distinct from overt brain injury.

Many common, acute complications of neonatal intensive care may compromise oxygen delivery and lead to transient hypoxemia including respiratory failure, sepsis, pulmonary hypertension, pneumothorax and patent ductus arteriosus.^12–14^ Isolated hypoxemia during preterm brain development is a significant risk factor for long-term neurodevelopmental disabilities.^15,16^ However, the role of isolated acute or chronic hypoxemia in learning and memory deficits has remained controversial given that preterm neonates frequently have co-morbid complications that preclude the identification of a well-defined cohort with hypoxemia alone.

We thus undertook mechanistic studies in preterm equivalent mice at P2 to determine if a clinically relevant exposure to mild hypoxia alone without ischemia is sufficient to disrupt hippocampal maturation without overt degeneration of hippocampal or cortical neurons associated with learning and memory. We developed a model that replicates the unique *ex-utero* physiology of the newborn brain as the neonate transitions to room air. P2 mice display brain development similar to preterm human infants at ∼26-28 weeks gestational age.^17,18^ Because mice are more resistant to hypoxia than humans,^19^ we employed well-tolerated conditions that resulted in transient tachycardia without bradycardia, moderate desaturations, no seizures, and vigorous feeding after hypoxia. We thus sought to achieve the equivalent of a brief, mild hypoxic insult in human preterm infants,^19^ since bradycardia, severe desaturations, seizures, and lethargy are often observed after moderate to severe hypoxia. In P3 mice, neonatal seizures, for example, are not reliably elicited without exposure to severe hypoxia.^20^ Here we report that a single mild hypoxic exposure was sufficient to trigger hippocampal memory impairment and persistently disrupt the maturation of hippocampal CA1 neurons without overt cerebral neuronal degeneration. Importantly, neuronal dysmaturation occurred independently of white matter injury, which supported intrinsic susceptibility of hippocampal neurons to hypoxia. We evaluated whether hypoxia affected long-term potentiation (LTP), a widely studied cellular mechanism of learning and memory. Neonatal hypoxia was sufficient to disrupt adult hippocampal CA3-CA1 synaptic strength, LTP and suppress calcium-sensitive synaptic SK2 channel activity, which is a key regulator of spike timing dependent neuroplasticity, including LTP. SK2 is a developmentally regulated Ca^2+^-activated K^+^ channel that acts as a negative feedback regulator^21,22^ to limit synaptic depolarization by AMPA and NMDA receptors. Hypoxia potently abolished synaptic SK2 function via a mechanism that did not involve reduced synaptic SK2 expression or redistribution within the synapse. Rather, loss of synaptic SK2 function was mediated by increased phosphorylation of SK2-associated calmodulin by the serine/threonine kinase CK2.

Our findings thus support a novel mechanism of long-term memory impairment in preterm survivors that results in neuronal dysmaturation rather than degeneration. Hippocampal CA1 neurons appear to be highly sensitive to a hypoxia-mediated mechanism that involves SK2 potassium channels that are a key regulator of developmental neuroplasticity and LTP strength.

## Materials and Methods

### Ethics statement

All studies were performed within the Oregon Health & Science University (OHSU) Department of Comparative Medicine using protocols approved by the OHSU Institutional Animal Care and Use Committee.

### Mouse hypoxia studies

At P2, litters of wild type C57/bl6 mice were reduced to 8 pups per litter. Prior to experimentation, adequate feeding was confirmed by visual confirmation of milk in stomach. P2 littermate mice were randomly assigned to 30 min exposure to heated room air (hereafter referred to as control, Con) or heated 8% O2 (hereafter referred to as hypoxia, Hx). Mice conditions were identified by limb tattoos and all pups were returned to the birthing dam. Maternal feeding and grooming behavior were confirmed after return to pups. Pups survived with dam for 24 or 72 hours for evaluation of acute injury responses or for 40-50 days for evaluation of all adult outcomes. Weaning was performed at P21. Sex was confirmed by visual inspection or genotyping. For all studies mice from at least 3 litters were represented in each group if possible.

### Physiological monitoring

Continuous heart rate (HR) and SpO_2_ were recorded from P2 pups using a neonatal probe and MouseOx pulse oximeter (Starr Life Sciences, PA) before, during and after 30 min Hx exposure. Recordings in pups of this age were prone to periods of poor data quality. All data was reviewed and periods of poor signal quality were removed. Mice of both sexes and exposures were weighed at multiple ages to monitor growth.

### Blood analysis

Before or at the end of the experimental treatment randomly selected mice underwent craniocervical dislocation. Mixed arteriovenous blood samples (30-50 mL) were immediately taken with heparinized capillary tubes from an incision piercing the carotid artery and jugular vein. Mice were decapitated immediately after. Metabolite content was analyzed using ABL725 blood gas analyzer (Radiometer Medical A/S).

### Mouse behavioral studies

Mice were habituated to the testing environment and handlers prior to behavioral testing. Hippocampal and amygdalar memory were tested using fear conditioning according to established protocols. Fear conditioning:^23^ Training for cued and contextual fear was conducted in conditioning chambers in sound-attenuated cubicles at P40-50. An 85 dB white noise cue (CS) was presented, followed by a mild 0.35 mA footshock (unconditioned stimulus). To assess contextual learning (hippocampus),^24^ 24 hours after training mice were returned to the same training apparatus and freezing, defined by cessation of all movement except respiration, and assessed for 5 min. To test for cued learning (amygdala and hippocampus),^23,25^ 48 hours after training mice were placed in an altered training apparatus (cleaned, novel scent, novel floor covering and novel exterior walls). Generalized freezing was assessed for 3 min, followed by two 3 min CS presentations with 3 min interval (ITI), and a 3 min post-CS period. The degree of freezing in the absence of CS (pre-CS, ITI, post-CS) defined baseline freezing and the freezing during CS presentations defined CS freezing. Training and testing sessions were recorded via a video camera (Wisecomm, Lithia Springs, GA) for analysis. Freezing was assessed by an observer blind to condition.

Sensorimotor performance was assessed on a rotarod as previously published.^26,27^ Briefly, mice were placed on an elevated rotating rod (diameter 3 cm; elevated 45 cm; Rotamex-5, Columbus Instruments, Columbus, OH), initially rotating at 5.0 rpm. The rod accelerated 1.0 rpm every 3 s. A line of photobeams beneath the rod recorded the latency to fall (seconds).

Each mouse received three trials per day, with no delay between trials, on three consecutive days

### Tissue collection and processing

For electrophysiology studies, mice were anesthetized using isofluorane and decapitated. Brains were immediately removed, hemisected, and placed into cold sucrose-aCSF (in mM: 80 NaCl, 2.5 KCl, 21.4 NaHCO_3_, 1.25 NaH_2_PO_4_, 0.5 CaCl2, 7.0 MgCl_2_, 20 glucose, 70 sucrose, 1.3 ascorbic acid; equilibrated with 95%O2/5%CO2).^21^ Transverse hippocampal slices (300 μm) were cut with a Leica VT1200S and transferred into a holding chamber containing regular aCSF (in mM: 125 NaCl, 2.5 KCl, 21.5 NaHCO3, 1.25 NaH_2_PO_4_, 2.0 CaCl_2_, 1.0 MgCl_2_, 12 glucose; equilibrated with 95%O_2_/5%CO_2_). Slices were incubated at 35°C for 30 min and then recovered at room temperature (22°C) for 1 hour before recordings were performed.

Tissues for immuno-histochemical analyses were initially immersed in 4% PFA in 0.1 M phosphate buffer (pH 7.4, 4°C) for 3 d, then stored in PBS with 0.05% sodium azide at 4°C until use. Tissues were sectioned coronally at 50 µm on a VT1000S and mounted on glass slides with VectaShield antifade (Vector Laboratories, Newark, CA) and coverslipped with 1.0 coverglass for standard microscopy or with ProlongGold antifade (ThermoFisher Scientific, Pittsburgh, PA) and topped with 1.5 coverglass for structured illumination microscopy.

For protein analysis, mouse brains were extracted, the hippocampus was micro-dissected and immediately frozen in liquid nitrogen for storage at -80°C.

### Golgi-Cox staining

Whole mouse brains were removed after decapitation and processed using a modified Golgi-Cox stain (https://cornell.flintbox.com/public/project/22082) produced by Drs. Lee and Jing of Cornell University (NY).^28^ All brain blocks were cut into 200 μm coronal slices using a vibrating microtome. All tissue slices containing the hippocampus were mounted on subbed slides, processed for Golgi visualization, dehydrated in a graded alcohol series, and topped with a 1.0 coverglass.

### Neuronal morphometry and spine counting

Well-filled Golgi-Cox stained pyramidal neurons within the CA1 hippocampus were identified. These neurons were traced using a 63x Plan Apochromat oil objective (NA 1.4, Carl Zeiss, Oberkochen, Baden-Württemberg, Germany) and reconstructed using a Leica DMREII inverted microscope and Neurolucida software (MBF Bioscience, Williston, VT). Criteria for tracing included: within CA1 hippocampus; complete filling of the cell body and dendrites; no visual obstruction by other filled objects; presence of a complete apical or basal dendritic compartment with minimal truncation (apical: 35/2812 [1.24%] ends; basal: 79/2581 [3.16%] ends). As previously described,^29^ NeuroExplorer software (MBF Bioscience) was used to generate morphometric analyses. Apical and basal dendritic arbors were analyzed independently. Sholl intersection (10 μm radii interval) and branch order analyses were performed to further assess dendritic complexity as previously described.^28^ Tracing and analyses were performed by persons blinded to condition.

### Dendritic spine counts

Neurolucida and NeuroExplorer software were used to quantify the density of the apical and basal dendritic spines on the same population of well Golgi-filled CA1 hippocampal neurons as previously described.^28^ Briefly, using a 63x oil objective (1.4NA), spine density was determined by counting all visible spines on 25 μm of terminal 3^rd^ order neurites initiating at between 100 and 200 μm from the cell soma.

### Immunohistochemistry

The detailed immunohistochemical protocols to visualize specific cell types were performed as previously described.^30–32^ Immunohistochemical procedures including antibodies and dilutions used are provided in Supporting Methods. Oligodendrocyte (OL) lineage cells were visualized with OL surface marker O4 in immature animals and OL transcription factor 2 (Olig2) in mature animals. Astroglia were visualized using glial fibrillary acidic protein (GFAP) or SRY-box transcription factor 9 (Sox9) antisera. Microglia and macrophages were visualized with an anti-ionized calcium-binding adapter molecule 1 (Iba1) antibody. Neurons were identified using a neuronal nuclear (NeuN) antibody. Axons were visualized with the pan-axonal neurofilament marker SMI-312. Anti-activated caspase-3-antibody (AC3) was used to identify apoptotic cells. Postsynaptic elements were visualized using postsynaptic density protein 95 (PSD95) antibody. Small conductance Ca^2+^-activated K^+^ channels (SK2) were identified with a rabbit polyclonal antibody as previously described.^33^Casein kinase 2 (CK2) mediated phosphorylation of calmodulin (CaM) was visualized with a phospho-calmodulin (Thr^79^, Ser^81^) polyclonal (pCaM) antibody.^34^ For fluorescent immunohistochemical studies, tissue sections were counterstained with Hoechst 33342.

### Unbiased quantification of cells

Sections immunostained for O4/AC3, Olig2/AC3, GFAP/Iba1, GFAP/Sox9, and NeuN counterstained with Hoescht 33342 were imaged using a Axio-Imager M2 upright microscope (Carl Zeiss) controlled by the StereoInvestigator software (RRID:SCR_002526, MBF Bioscience) and Hamamatsu C11440 (Hamamatsu, Hamamatsu City, Japan) digital camera. At least 2 sections from the mid hippocampal level were outlined and ROI defining the entirety of cerebral cortex, white matter, and hippocampi were drawn using StereoInvestigator. Using the Optical Fractionator tool and stereological unbiased counting, at least 10 sites were defined by systematic random sample placement with guard zones and counting grid. Nuclei associated with a label of interest were counted using a 20x or 40x air objective as appropriate based on density. Degenerating O4-positive cells were confirmed to contain degenerating pyknotic nuclei.

### Image quantification

Unbiased semi-quantitative analyses of immunohistochemical markers were determined by a blinded individual as previously described.^35,36^ Briefly, images of staining for GFAP, Iba1 or pCaM in the medial primary somatosensory cortex, CA1 hippocampus, and cingulate/corpus callosal white matter were acquired in the coronal plane at the mid-hippocampal level using a 20x objective with fixed image acquisition settings by a blinded observer on Axio-Imager M2 upright microscope with a Hamamatsu C11440 using HCI Image (Hamamatsu). Myelin basic protein (MBP) staining was photographed using a 10x objective in two adjacent regions at the mid hippocampal region comprising the midline to the lateral extent of the hippocampus using identical acquisition settings. Camera acquisition settings were optimized for each label to improve signal:noise ratios by maximizing occupation of the image histogram while having no under-or overexposed pixels in true negative or positive staining. Artifactual staining, if present, was removed during image segmentation or as otherwise noted. For GFAP and Iba1, images were segmented into cortical, white matter, and hippocampal regions of interest (ROI). Because pCaM staining showed clear layer specific staining, hippocampal images were segmented into stratum oriens, pyramidal, radiatum, and lacunosum moleculare; and cortical images into neuronal layers II-VI. A pixel-intensity histogram was generated for each ROI using ImageJ (NIH; rsbweb.nih.gov/ij/index.html) and exported to a spreadsheet. The peak of the histogram was calculated using the three highest frequency bins, and the histogram curve integrated towards the background pixel side of the peak and a value obtained for this region’s area. This area was then doubled to estimate the total distribution of background voxels in the image. The total background area was subtracted from the total ROI area to define the labeled area.

### Hippocampal electrophysiology

For hippocampal field recordings, slices were placed on a temperature controlled (32°C) chamber infused continuously with oxygenated aCSF at 1.5 mL/min. Slices were visualized using a fixed-stage, upright microscope (Zeiss Axioskop 2FS, Carl Zeiss) with a 5x objective. Electrodes were pulled from borosilicate pipettes (Sutter Instruments) with resistances of 2 MW when used for recording or 1 MW when used for stimulation and were filled with aCSF. The stimulating electrode was placed in the CA1 stratum radiatum 100-200 µm from stratum pyramidal and the recording electrode laterally separated by 100-300 µm equidistant from the stratum pyramidal. CA3 Schaffer collateral axons were stimulated using a Digitimer constant current stimulus isolation unit (AutoMate Scientific). Stimulus duration was 0.1 ms allowing for clear separation of fiber volley (FV) from the preceding stimulus artifact. CA1 field excitatory post-synaptic potential (fEPSP) recordings were obtained using a Multiclamp 700B (Molecular Devices, Sunnyvale, CA) interfaced to an ITC-18 analog-to-digital converter (HEKA, Harvard Bioscience Inc, Holliston, MA). Signals were filtered at 5 kHz, digitized at 20 kHz, and transferred to a computer using Patchmaster software (HEKA, Harvard Bioscience Inc, Holliston, MA). Field responses were elicited at 0.05 Hz. All experiments were performed in the presence of SR95531 (5 mM, Tocris Bio-Techne, Minneapolis, MN) and CGP55845 (2.5 mM Tocris) to block inhibitory synaptic transmission. The CA1:CA3 axons were severed to eliminate recurrent excitation within the CA3 subfield. To measure synaptic input output relationship of fEPSP (I/O), a series of subthreshold stimuli were delivered at 20 µA intervals from 20 µA to 100 µA. The derivative (dV/dT) of the initial 2–3 ms onset of the fEPSP was measured (fEPSP slope).

The slope of a line fit to the fEPSP slope:FV relationship reflects the input output relation of CA3:CA1 synaptic transmission. Paired-pulse responses were recorded using a 50 ms interpulse interval (20 Hz) and expressed as a ratio of the second pulse over the first pulse (P2:P1). Prior to the induction of long-term potentiation (LTP) at CA3:CA1 synapses, the stimulus intensity was set at 0.25 of the minimum stimulus that produced backpropagating action potentials (pop spikes) to avoid additional spike timing dependent plasticity during LTP recordings. A 10 min stable baseline period was established before delivering a theta burst stimulation (TB) train [a single burst consists of five fEPSP stimuli delivered at 100 Hz and ten bursts delivered at 5 Hz per sweep, three TB sweeps were delivered at 30 s intervals]. Following TB, the fEPSP was recorded for 60 min. The fEPSP slope was analyzed for individual slices from 50–60 min post-TB and normalized to baseline before LTP induction. Diary plots of LTP were obtained by binning all slices for each treatment at 1 min intervals.

Whole-cell patch-clamp recordings were obtained using a Multiclamp 700B amplifier (Molecular Devices). Signals were filtered at 5 kHz, digitized at 20 kHz, and transferred to a computer using an ITC-18 and Patchmaster. Recordings were obtained at room temperature (22°C). CA1 pyramidal cells were visualized with infrared–differential interference contrast optics (Zeiss Axioskop 2FS, Carl Zeiss). Electrodes pulled from borosilicate pipettes (Sutter Instruments) had tip resistances of 3 MW for recording and 2 MW for stimulation. Patch pipettes for EPSP recordings were filled with a K-gluconate internal solution containing (in mM)^37^ 133 K-gluconate, 4 KCl, 4 NaCl, 2 MgCl 2, 10 HEPES, 4 MgATP, 0.3 Na 3GTP, 10 K-phospho-creatine (pH 7.3).

EPSPs were recorded in whole-cell current-clamp mode and voltages were not corrected for a junction potential of -17 mV. All recordings used cells with a resting membrane potential less than -50 mV and a stable input resistance that did not change by more than 20%. Cells were biased to -65 mV and the input resistance was determined from a 25 pA hyperpolarizing current injection pulse given 500 ms after each synaptically evoked EPSP. The stimulating electrode was placed ∼150 μm into and 50-100 μm lateral to the apical dendritic field and placement was adjusted to obtain monosynaptic EPSPs. To assay for SK2 channel activity, apamin (100 µM) was added to the bath solution, the baseline was recorded for 5-10 min and EPSPs were recorded for another 20 min. To evaluate the contribution of casein kinase 2 (CK2) to SK2 activity, the selective cell-permeable CK2 inhibitor 4,5,6,7-Tetrabromobenzotriazole (TBB, 10 µm) was added to the bath for 20 min prior to baseline determination in select experiments. Data were analyzed in Igor Pro (Wavemetrics, Lake Oswego, OR).

### Western Blotting

Analysis was performed once all frozen samples for a particular experimental group were collected. To avoid batch effects, we prepared protein lysates in NE-PER buffer (ThermoFisher Scientific) supplemented with protease and phosphatase inhibitors from all the samples on the same day. Protein concentrations were estimated by the standard BCA assay (ThermoFisher Scientific), and 35 μg of total protein lysate was employed for all downstream western blotting assays. All protein lysates were subjected to SDS-PAGE using 4-20% gels (Bio-Rad Laboratories) and blotted onto Immobilon-FL PVDF membrane (Millipore Sigma). Primary and secondary antibodies used are summarized in Supporting Methods and include GluA1, GluA2, GluN2A, GluN2B. To quantify changes in signal intensities for all the blotted proteins, we normalized the densitometry values to b-actin. Densitometry values were obtained via Image Studio Lite (LI-COR Biosciences).

### Structured illumination microscopy (SIM)

We obtained 3D image stacks from CA1 stratum radiatum between 100 and 300 µm from the pyramidal cell layer (Fig. 6D). Three samples were taken in the mediolateral axis from the mid-hippocampal level. Image data was detected using dual PCO edge sCMOS cameras on a Zeiss Elyra7 Structured Illumination Microscope using a 63x1.4 NA lens. Samples were illuminated sequentially with a 488 nm or 561 nm laser line split at 560 nm and emission detected through a 495-550 nm and 570-620 nm filters, respectively. Low magnification was used to locate the stratum radiatum of CA1 using nuclear staining as a guide and z-stacks of 10-20 µm range at 0.11 µm steps were acquired in three randomly selected positions per hemisphere. Care was taken to minimize blood vessels in the region of interest. Post-processing with SIM^2^ algorithm (Zen Black 3.0) resulted in images with 30 nm pixels in the XY dimension. To account for chromatic shift, a bead slide of 0.1 µm TetraSpeck beads (T7279) mounted identically to the sample slides was imaged at the end of each microscope session. Channel alignment matrices calculated in Zen Black were used to align image stacks prior to further analysis. The microscope operator was blinded to experimental conditions.

### SIM Image processing and quantification

SIM image analysis is adapted from previously described methods.^38,39^ Aligned and SIM^2^ processed images were analyzed in Imaris v9.9 (Andor Technologies Inc) to identify PSD95 puncta (488 channel) and SK2 puncta (555 channel).^38^ Puncta were identified using the ‘Spots’ feature in Imaris with parameter settings determined from several randomly selected images. To identify puncta with real signal, thresholds were calculated to eliminate 95% of puncta detected from samples treated with only secondary antibodies. Autofluorescence of blood vessels in the 488 channel was identified using Imaris machine learning algorithm and masked out using the ‘Surfaces’ feature prior to further analysis. Blood vessel-masked images were analyzed to find the distance between PSD95 puncta to the nearest SK2 puncta. Images were batched processed using the pre-defined settings. Analysis was performed blinded. Post-processing with SIM^2^ algorithm (Zen Black 3.0) resulted in images with up to 60 nm resolution in the XY dimension and 200 nm in the Z dimension. Each image underwent frequency distribution (bin width of 10 nm) and was fit with a sum of two Gaussian distribution. The near curve characteristics relate to colocalized puncta and were utilized for further analysis either as a fraction of total or normalized to image volume as indicated.

### Statistical Analysis

All images and data were acquired, and data analyzed blinded to condition. Analyses were performed using a two-sided alpha 0.05 and beta 0.2 unless otherwise noted. Data were tested for normal distribution using the D’Agostino & Pearson test and parametric vs nonparametric tests chosen as appropriate. Sex was analyzed as a biological variable by 2 way ANOVA unless otherwise noted. If sex effects were not present, sexes were pooled to increase statistical power. Two group comparisons were performed by either student’s t-test or Mann-Whitney as appropriate. Multiple group comparisons were performed with either ANOVA or Kruskal-Wallis as appropriate followed by the Dunn post-hoc correction. Growth data was log transformed, fit with an exponential growth curve and within-sex comparisons made with extra sum-of-squares F test. Correlation analysis was performed using a Spearman correlation due to non-normal distribution followed by two-stage linear step-up procedure of Benjamini, Krieger and Yekutieli, * = Q: 1%. To reduce type 1 and type 2 errors,^40^ linear mixed model (LMM) approach was utilized for analyses with nested subject data, missing data, and multiple distribution types such as behavioral data, neuron structure data, spine density data, multiple sample cell counts and electrophysiology data. Unless otherwise noted post hoc comparisons were made using Sidak. For Sholl and branch order analyses, a LMM was performed at each radius or order followed by two-stage linear step-up procedure of Benjamini, Krieger and Yekutieli, * = Q: 1% due to the large number of comparisons. For LMM models, sex was included in all models, regardless of significance. If applicable, analysis for location and observer were included in initial models and removed if non-significant. LMM data is reported as estimated marginal mean (EMM) with standard error (SE).

## Results

### Mild hypoxia is well tolerated in neonatal mice

We first established conditions that would be physiologically well tolerated and would minimize the risk that neonatal hypoxia exposure might alter maternal rearing and pup behavior. Preterm equivalent P2 mice were removed from the home cage and randomly assigned to a 30 min exposure to humidified room air (RA) or 8% inspired oxygen (FiO_2_; Hx; Fig. 1A). Because mice are more resistant to hypoxia than human, we first tested these conditions, which we previously found produce no neuronal degeneration in mice even in the setting of ischemia.^41^ To determine if P2 mice exposed to 8% FiO_2_ sustained cardiovascular instability, as previously reported at 5.7%,^11^ we measured heart rate (HR) and oxygen saturations before, during, and after hypoxia (Fig. 1B, C). Within the first 20 sec of exposure to 8% FiO_2_, mice had a significant drop in oxygen saturations that reached a mean of 84.3 ± 3.9% (mean ± SD) by 3 min of exposure. This degree of hypoxia was not accompanied by a drop in heart rate.^11^ Rather mild tachycardia was observed that increased from 575 ± 15 to 665 ± 67 beats per minute (Fig. 1B, mean ± SD). Recovery to baseline heart rate and normal oxygen saturations occurred over 45 sec after return to room air. No abnormal behavior, movement, seizures, or evidence of cardiovascular instability was observed in any Hx exposed or sham room air pups (hereafter referred to as control; Con). After animals returned to the dam in the home cage, maternal nursing was confirmed by visualization of milk in the pups’ stomachs. Subsequent weight gain did not differ between Hx and Con, but typical sexual dimorphism was noted over the experimental period (Fig. 1D). A normal baseline glucose for age of 5.1 ± 2.1 mmol/L was also confirmed^42^ and did not significantly differ after 30 min of Con (5.1 ± 1.6) vs Hx (3.1 ± 1.2 mmol/L; n = 8 per group, ANOVA; Tukey post hoc correction, mean ± SD) conditions (Fig. 1E). Thus, a single 30-minute exposure to 8% FiO_2_ generated mild hypoxia without bradycardia that mimicked key features of hypoxemia without associated adverse physiologic or metabolic events, similar to that frequently encountered by preterm neonates under similar conditions.^43,44^

**Figure 1:**
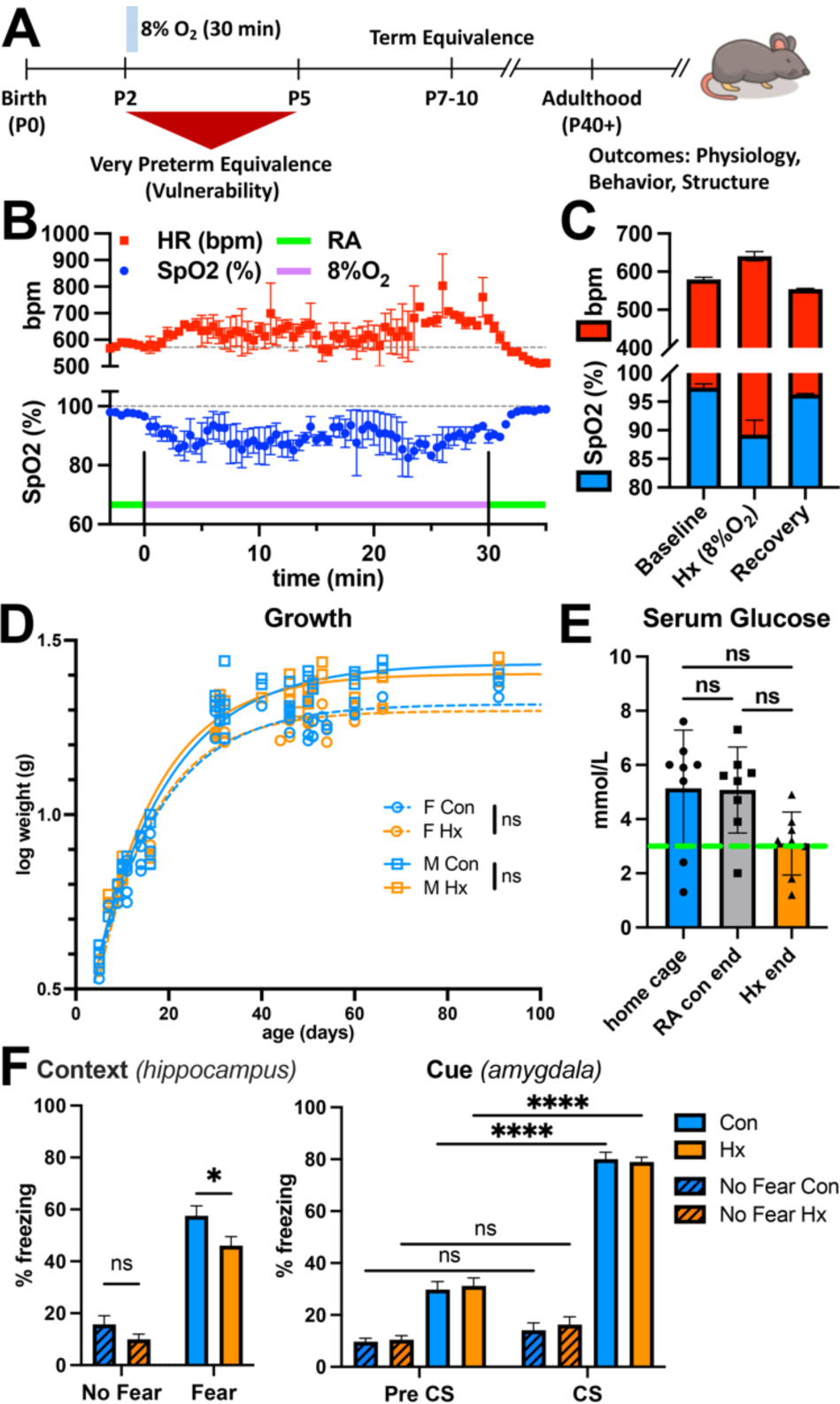
Responses to neonatal Hx (8%O_2_; 30-min) were mild and caused persistent hippocampal memory deficits in mice. *A*) Schematic of experimental approach for murine neonatal hypoxia protocol relative to human brain development. *B, C*) Neonatal Hx was associated with transient tachycardia and moderate SpO_2_ desaturations. *D*) Growth was not impaired after neonatal Hx in male (m) or female (f) mice. Log normalized data fit with exponential plateau showed no effect of Hx on growth (f Con n = 75 vs. Hx n = 57; m Con n= 72 vs. Hx n = 76; extra sum-of-squares F test). *E*) 30 min maternal separation did not cause hypoglycemia in Con or Hx exposed mice (n = 8/group; mean ± SD; one-way Anova with Tukey) and closely approximated previously published P3 control data^42^ (*green dashed line*). *F*) Neonatal Hx caused hippocampal-based memory deficits in adult mice determined with a fear-conditioning test. Hx mice that were fear conditioned (n = 27) displayed a significant reduction in % freezing duration at 6-8 weeks (n = 30 Con; Welch’s t-test). Consistent with a Hx-mediated hippocampal specific memory deficit, a cue test of amygdala function found testing was effective, but there was no significant effect of Hx during pre-conditioning (pre-CS) or during a conditioned stimulus (CS; 2-way ANOVA with post-hoc Dunn, mean ± SEM). * p<0.05, **** p<0.0001.

### Hippocampal learning deficits occur in adult mice exposed to mild neonatal Hx

To determine if neonatal Hx is sufficient to disrupt adult hippocampal learning and memory, we evaluated hippocampus and amygdala-dependent learning through fear conditioning at P40-50. During training, animals were exposed to an aversive foot shock that was associated with a unique chamber and auditory cue. Contextual learning is hippocampal and was assessed by returning mice to the same chamber 24 hours after training to assay for freezing behavior. Cued learning is primarily mediated by the amygdala and was assessed by returning mice to a new chamber 48 hours after training to assay for freezing behavior before and after exposure to the auditory cue.^23^ Contextual freezing behavior was significantly reduced in adult animals exposed to neonatal mild Hx vs littermate controls (Fig. 1F). Sex was analyzed as a variable and was not found to be significant or interact with Hx exposure. A robust freezing response to the auditory cue was observed that did not differ between Hx and con animals (Fig. 1F). Thus, hippocampal, but not amygdala-mediated learning was impaired in adult animals of both sexes after exposure to a single mild episode of neonatal Hx.

### Hippocampal CA1 dendritic morphology in adult mice is disrupted after neonatal Hx

We next determined if lasting hippocampal memory deficits were accompanied by disturbances in neuronal maturation. The morphology of adult (P50) hippocampal CA1 neurons was visualized utilizing the Golgi-Cox staining method. Apical and basal dendritic arbors were traced and quantified in 3D using Neurolucida and Neuroexplorer (MBF), respectively. Figure 2A shows the results of a Sholl analysis of dendritic arbor complexity of young adult CA1 hippocampal neurons after P2 Hx (Con: n = 75 neurons, 17 animals, 9 m, 8 f; Hx: n = 135 neurons, 21 animals, 10 m, 11 f). Neuronal morphology was analyzed using a Linear Mixed Model approach, in order to minimize the risk of type 1 or type 2 errors that are prevalent with common statistical approaches to neuronal morphology data.^40^ Analyses within all groups were performed independently by two observers. Initial statistical models included observer as well as sex and condition as variables, however observer was not found to contribute to effect in any output and was dropped from the final models. Sex was not a significant factor in the majority of outputs, unless otherwise noted. Sholl analysis revealed that maturational responses to Hx differed between basal and apical dendrites (Fig. 2A, B). CA1 basal neurites are oriented toward the stratum oriens (s.o.), while apical dendrites interface with CA Schaffer collaterals within the stratum radiatum (s.r.) and extend into the stratum lacunosum moleculare (s.l.m.). Hx mediated disturbances were most prevalent in CA1 basal dendritic arbors, which showed increased intersections across much of the tree. Because CA1 dendritic morphology can vary between sexes,^45^ we evaluated for sex dependence. Neurons from female controls displayed a trend towards more primary dendrites per neuron but less branching per dendrite (Table 1), which resulted in no sex difference in total dendritic length (estimated marginal mean ± standard error: EMM ± SE; Con, 577 ± 38 µm, Hx 533 ± 47 µm). The increase in primary dendrites after Hx appeared to be driven primarily by male animals (f, Con 4.02 ± 0.19, Hx 3.89 ± 0.14; m, Con 3.10 ± 0.23, Hx 3.98 ± 0.18, interaction p = 0.006, EMM ± SE). No other variables showed a significant interaction between Hx treatment and sex (Table 1, Fig. 2C). Because there was a significant effect of Hx exposure and sex on the number of basal primary dendrites, additional morphological parameters were normalized per dendrite (Table 1, Fig. 2C). This was not required on apical analyses as CA1 neurons always have a single primary apical dendrite. Notably, the Hx-related increase in basal complexity was due to an increase in both the number of primary dendrites as well as increased dendritic branching reflected by an increase in nodes, ends and length per dendrite (Table 1, Fig. 2C). This resulted in a 74% increase in total basal dendritic length in Hx exposed animals (Con 405 ± 48 µm; Hx 706 ± 36 µm, EMM ± SE). Surprisingly, altered dendritic maturation was not present within the apical dendritic arbor (Table 1), which extends through the s.r. and s.l.m. Apical dendritic length, nodes, ends and Sholl analyses all showed a trend toward increased complexity but no measures reached statistical significance. In order to confirm if apical and basal compartments have different morphological responses to Hx, correlational analyses between, within and across apical and basal arbors was performed. Correlational analysis confirmed that factors that drive complexity within either the basal or apical dendritic arbors were highly correlated (Fig. 2D, blue), however, no significant correlation or anticorrelation exists between the two regions (Fig. 2D, white). Thus, persistent morphometric disturbances in adult hippocampal CA1 neuronal maturation occurred after neonatal Hx, which was neuronal subregion specific.

**Figure 2:**
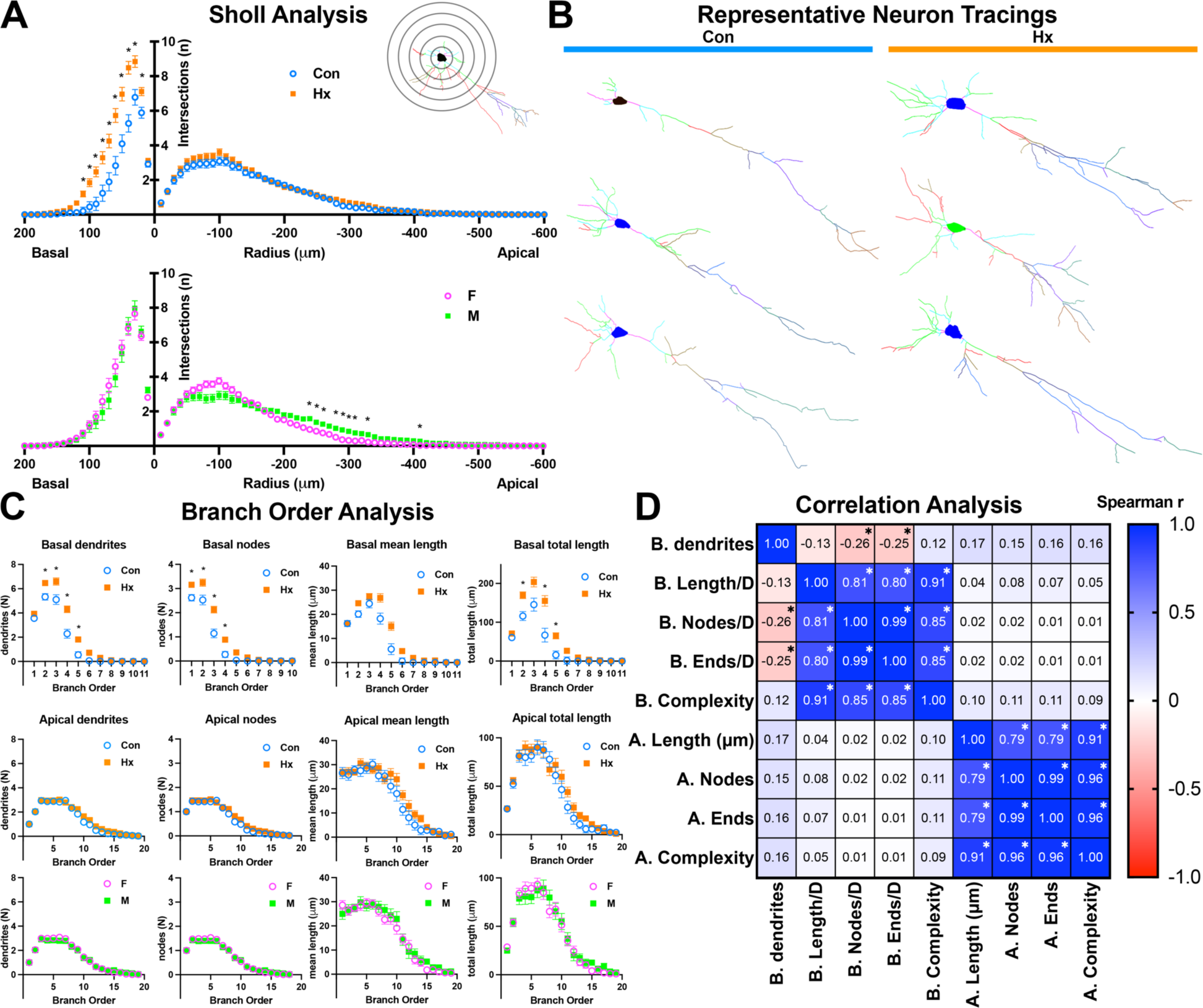
Neonatal Hx disrupts **adult** CA1 neuron dendritic complexity. *A*) Neuronal complexity determined by Sholl intersection analysis (*schematic inset image*) was increased after Hx across much of the basal dendritic tree. Analysis of sex revealed minor differences in the apical arbor. (n = Hx 131 neurons, 21 mice, 10 m, 11 f; Con 72 neurons, 15 mice, 7 m, 8 f; LMM at each radius, EMM ± SEM). *B*) Representative tracings of Con and Hx exposed CA1 neurons. *C*) Branch order analysis confirmed Hx increased basal neurite number, branching, and total length (same data and method as above). *D*) A correlation matrix shows that factors driving basal and apical complexity do not correlate (*white*), but do within each arbor (*blue*, majority of data did not pass normality test, thus a Spearman correlation was used). * indicates significance after multiple comparison correction with two-stage linear step-up procedure of Benjamini, Krieger and Yekutieli, Q: 1%.

**Table 1.** Basal Dendritic Tree.

| Variable |  | Significance (p) |  | Sex<sup>†</sup> | Mean<sup>‡</sup> | 95% CI |
| --- | --- | --- | --- | --- | --- | --- |
| # 1° dendrites | treatment | 0.048* | Con | f | 4.02 | 3.65–4.40 |
|  | sex | 0.027* |  | m | 3.10 | 2.66–3.54 |
|  | interaction | 0.006** | Hx | f | 3.88 | 3.61–4.15 |
|  |  |  |  | m | 3.98 | 3.63–4.33 |
| nodes/dendrite | treatment | <0.001*** | Con | f | 1.71 | 1.27–2.16 |
|  | sex | 0.014* |  | m | 2.23 | 1.69–2.75 |
|  | interaction | 0.846 | Hx | f | 2.42 | 2.10–2.74 |
|  |  |  |  | m | 3.00 | 2.59–3.42 |
| ends/dendrite | treatment | <0.001*** | Con | f | 2.76 | 2.31–3.21 |
|  | sex | 0.010* |  | m | 3.25 | 2.71–3.78 |
|  | interaction | 0.695 | Hx | f | 3.47 | 3.15–3.79 |
|  |  |  |  | m | 4.14 | 3.72–4.55 |
| length/dendrite | treatment | <0.001*** | Con |  | 118 µm | 92–144 µm |
|  | sex | 0.882 | Hx |  | 188 µm | 168–208 µm |
|  | interaction | 0.730 |  |  |  |  |
| complexity<sup>f</sup> | treatment | 0.002** | Con |  | 3972 | -423–8366 |
|  | sex | 0.206 | Hx |  | 12938 | 9619–16257 |
|  | interaction | 0.177 |  |  |  |  |

**Table 1:** P2 mild Hx results in altered adult CA1 hippocampal morphology. (con: n = 75 neurons, 17 animals, 9 m, 8 f; hx: n = 135 neurons, 21 animals, 10 m, 11 f), Linear Mixed Model, \* p < 0.05, \*\* p < 0.01, \*\*\* p < 0.001.
| Variable |  | Significance (p) |  |  | Mean<sup>‡</sup> | 95% CI |
| --- | --- | --- | --- | --- | --- | --- |
| nodes | treatment | 0.071 | Con |  | 12.1 | 10.9–13.2 |
|  | sex | 0.719 | Hx |  | 13.4 | 12.5–14.3 |
|  | interaction | 0.475 |  |  |  |  |
| ends | treatment | 0.088 | Con |  | 13.3 | 12.19–14.49 |
|  | sex | 0.622 | Hx |  | 14.6 | 13.71–15.53 |
|  | interaction | 0.424 |  |  |  |  |
| length | treatment | 0.073 | Con |  | 768 µm | 696–839 µm |
|  | sex | 0.791 | Hx |  | 851 µm | 795–908 µm |
|  | interaction | 0.515 |  |  |  |  |
| complexity<sup>f</sup> | treatment | 0.094 | Con |  | 82457 | 62209–102705 |
|  | sex | 0.970 | Hx |  | 104455 | 88481–120428 |
|  | interaction | 0.460 |  |  |  |  |

†: sex-based means are displayed for variables that showed significant sex-dependence
‡: estimated marginal means are shown
f: complexity = [sum of the terminal orders + number of terminals] \* [total dendritic length / number of primary dendrites]

### Early or delayed white matter injury (WMI), cell death and inflammation do not accompany neonatal Hx

We next evaluated the contribution of neonatal WMI, cell death and inflammation to impaired hippocampal learning after Hx. Early and delayed WMI are the primary forms of overt injury seen in preterm survivors and related animal models.^31,32,35,46^ Late oligodendrocyte progenitors (preOLs) are the primary cell type that degenerates in neonatal WMI and is accompanied by glial reaction involving astrocytes and microglia.^31^ We analyzed injury responses in the white matter, hippocampus and cerebral cortex given their overlapping roles in distinct aspects of learning and memory. We quantified intact and degenerating O4 antibody labeled preOLs at 24 h (P3, Fig. 3A) and 72 h (P5, Fig. 3B) after Hx at P2. No change in total oligodendrocyte density, degenerating OL lineage cells (O4^+^ + pyknotic nuclei) or OL lineage cell apoptosis (O4^+^ + activated caspase-3, AC3) was identified at 24 or 72 h in any region. Our findings support that neonatal Hx does not cause acute loss of OL lineage cells, degeneration of neurons or other glia in cortical or hippocampal gray matter, or in corpus callosum white matter at 24 or 72 h after Hx at P2.

**Figure 3:**
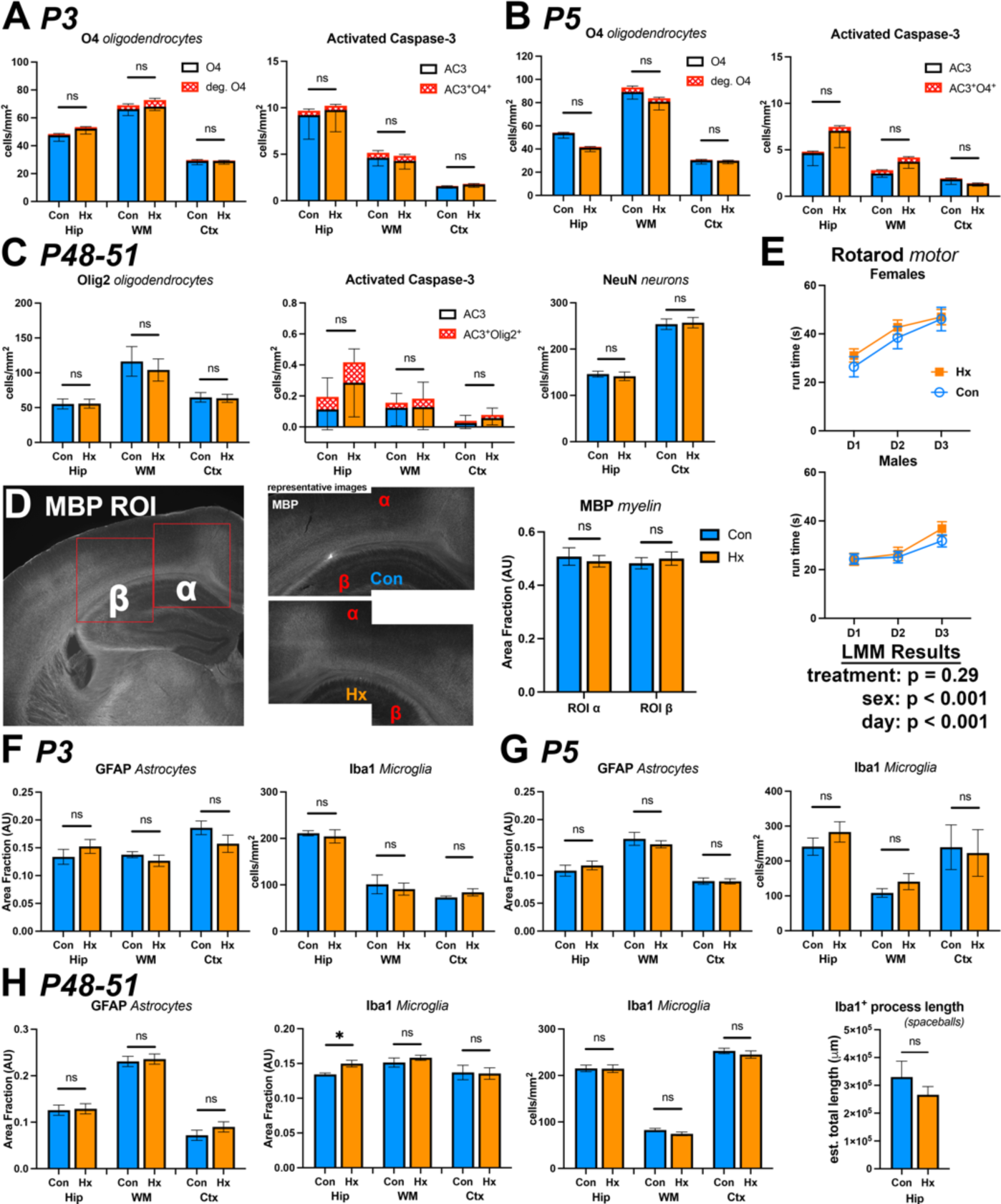
Oligodendrocye (OL) lineage maturation and myelination are not altered after neonatal Hx in white or gray matter structures. No change in OL density (O4+), OL death (O4+ pyknotic nuclei), or apoptotic cell death (activated caspase-3, AC3) were observed in white matter (WM), hippocampus (Hip), or cortex (Ctx) at 24 hours (*A*, P3) or 72 hours (*B*, P5) after Hx (n = 24 images, 8 mixed sex mice/group). *C*) No Hx-induced change in OL (Olig2) density, OL apoptosis, or overall apoptosis (AC3) was identified. Neuron density within gray matter structures was not altered after neonatal Hx. *D*) Evaluation of myelination in adult mice with myelin basic protein (MBP) staining in adjacent fields of the corona radiata (α, β) did not reveal any evidence of dysmyelination. *E*) Rotarod motor testing was not altered after Hx, but did reveal expected sex and training day differences (n = Con 30 mice, F 6, M 24; Hx 32 mice, 14 f, 17 m; EMM ± SEM; LMM). Neonatal Hx does not cause early or late astrocytic (GFAP) or microglial (Iba1) inflammatory responses. Acute (*F*) and subacute (*G*) glial reactivity were not found (n = 8 mixed sex per group, mean ± SEM). *H*) Hx exposure did not alter adult astrocyte and microglial markers (n=8 mixed sex per group, mean ± SEM). Iba1 area fraction in the hippocampus showed a small significant increase that was not confirmed by unbiased cell count or process length (spaceballs) measures.

To determine if early inflammation occurs after neonatal Hx, we analyzed subacute astrocyte and microglial activation within the hippocampus, white matter and cortex (Fig. 3F, G). Astrocytes (GFAP^+^) and microglial (Iba1^+^) showed no increase in staining or change in staining pattern 24 or 72 h after Hx that would indicate early glial activation. Quantification of microglial reactivity found no alterations in Iba1 area fraction or density after P2 Hx. In summary, we found no evidence of acute oligodendrocyte injury, cell death or apoptosis, or inflammatory astrocyte or microglial responses to P2 Hx in the WM, hippocampus, or cortex.

To determine if neonatal Hx triggered delayed WMI that contributes to impaired hippocampal function, dysmyelination and delayed cell death were evaluated in young adult animals (P40-50; Fig. 3C). Myelin integrity was evaluated within the corpus callosum, corona radiata, and cerebral cortex by MBP staining at P50 (Fig. 3D). Estimation of MBP staining within the corona radiata, corpus callosum, and adjacent cortex metric showed no reductions in MBP consistent with dysmyelination after Hx (Fig. 3D, graph). Neurofilament staining with SMI 312 similarly showed no evidence of axonal injury, including no focal swellings, altered staining intensity, or fragmentation within the hippocampus, WM or corona radiata (Sup. Fig. 1). In addition, adult rotarod testing was used to evaluate for functional motor deficits. Rotarod testing showed that females performed better than males across multiple days of learning, but no effect of Hx exposure for same-sex littermates (Fig. 3E).

Altered OL lineage maturation or accumulation of immature OLs has been reported in multiple perinatal hypoxia models.^31,32,46^ However, OL lineage (Olig2) density was not altered after neonatal Hx in young adult animals in the WM, cortex, or hippocampus (Fig. 3C, Olig2). To broadly evaluate for delayed programmed cell death, AC3 was quantified. No change in the low baseline rate of cell death within WM, cortex, or hippocampus was observed (Fig 3C, activated caspase-3). Similarly, no increase in OL lineage apoptosis was observed by double labeling for AC3 (Fig 3C, AC3^+^Olig2^+^). To determine if neonatal Hx contributed to hippocampal dysmaturation through delayed neuronal cell death or aberrant neurogenesis, neuronal responses to P2 Hx at P50 were also evaluated. Analysis of NeuN^+^ neurons in the cortex and hippocampus found no changes in neuronal density after Hx (Fig. 3C, NeuN).

Since delayed astrocyte or microglial activation has been shown to contribute to injury progression in several neonatal models,^35,47,48^ we quantified GFAP, Sox9 and Iba1 staining at P50 from P2 Hx vs Con littermates (Fig. 3H). At P50, GFAP staining was not increased in any region (Fig. 3H, GFAP), and no effect on Sox9^+^ cell density was observed (hippocampus, Con 622 ± 39, Hx 699 ± 39, p = 0.18; white matter, Con 346 ± 28, Hx 323 ± 27, p = 0.56; cortex, Con 677 ± 41, Hx 641 ± 35, p = 0.50, n = 16 con [8f, 8m], 14 Hx [6f, 8 m], two sided t test after no sex effect on 2 way ANOVA, mean ± SEM), consistent with no chronic astrocyte reactivity due to Hx. There was a minimal but statistically significant increase in Iba1 staining by area fraction measurement in the P50 hippocampus that was not accompanied by an increase in microglial number or altered staining pattern (Fig. 3H, Iba1). The spaceballs probe, an unbiased stereological assessment of process morphology was used to evaluate if a broad increase in microglial reactivity without increased cell density might be present (Fig. 3H, Iba1 process length). Hx exposure did not result in altered process morphology. Thus, no consistent change in microglial morphology or number was found to support chronic microglial activation after neonatal Hx.

In summary, our evaluation of multiple timepoints, cell types, and injury markers in the hippocampus, cerebral cortex and callosal WM found no evidence of acute or delayed cell death, inflammation or dysmyelination or that might be related to hippocampal neuronal dysmaturation or altered hippocampal learning in young adult mice exposed to mild Hx at P2.

### Hippocampal CA1 cellular learning and memory mechanisms are disrupted by neonatal Hx

The primary driver of hippocampal CA1 action potential output is CA3 Shaffer collateral synapses onto CA1 apical dendrites (Fig. 4A). These synapses undergo lasting activity dependent strengthening, or long-term potentiation (LTP), which is thought to be the primary cellular correlate of hippocampal learning and memory.^49,50^ The fast phase of LTP is primarily due to trafficking of AMPAR into the postsynaptic density, where they promote enhanced excitatory postsynaptic potentials (EPSPs) in response to glutamate release.^51–53^ To determine if Hx disrupts LTP, we determined the percent LTP induction at CA3-CA1 synapses in young adult mice, which were P2 Hx or Con littermates. Extracellular field EPSPs (fEPSPs, Fig. 4B) were induced at 25% of the intensity required to initiate pop spikes (backpropagating action potential) in each given slice at 20s intervals. After a 10 min stable baseline was recorded, LTP was induced using a theta burst (TB) protocol^54^ and fEPSP responses were measured for 60 min after LTP induction (Fig. 4C diary plot). LTP was measured from the average of 15 traces preceding TB induction and 55-60 min after TB. The Hx group (n = 23 slices, 12 mice) displayed a significant decrease in LTP vs. controls (n = 14 slices, 6 mice; Fig 4C scatterplot). Thus, neonatal Hx resulted in persistently altered information coding within the hippocampus CA3-CA1 circuit that may contribute to altered hippocampal learning and memory.

**Figure 4:**
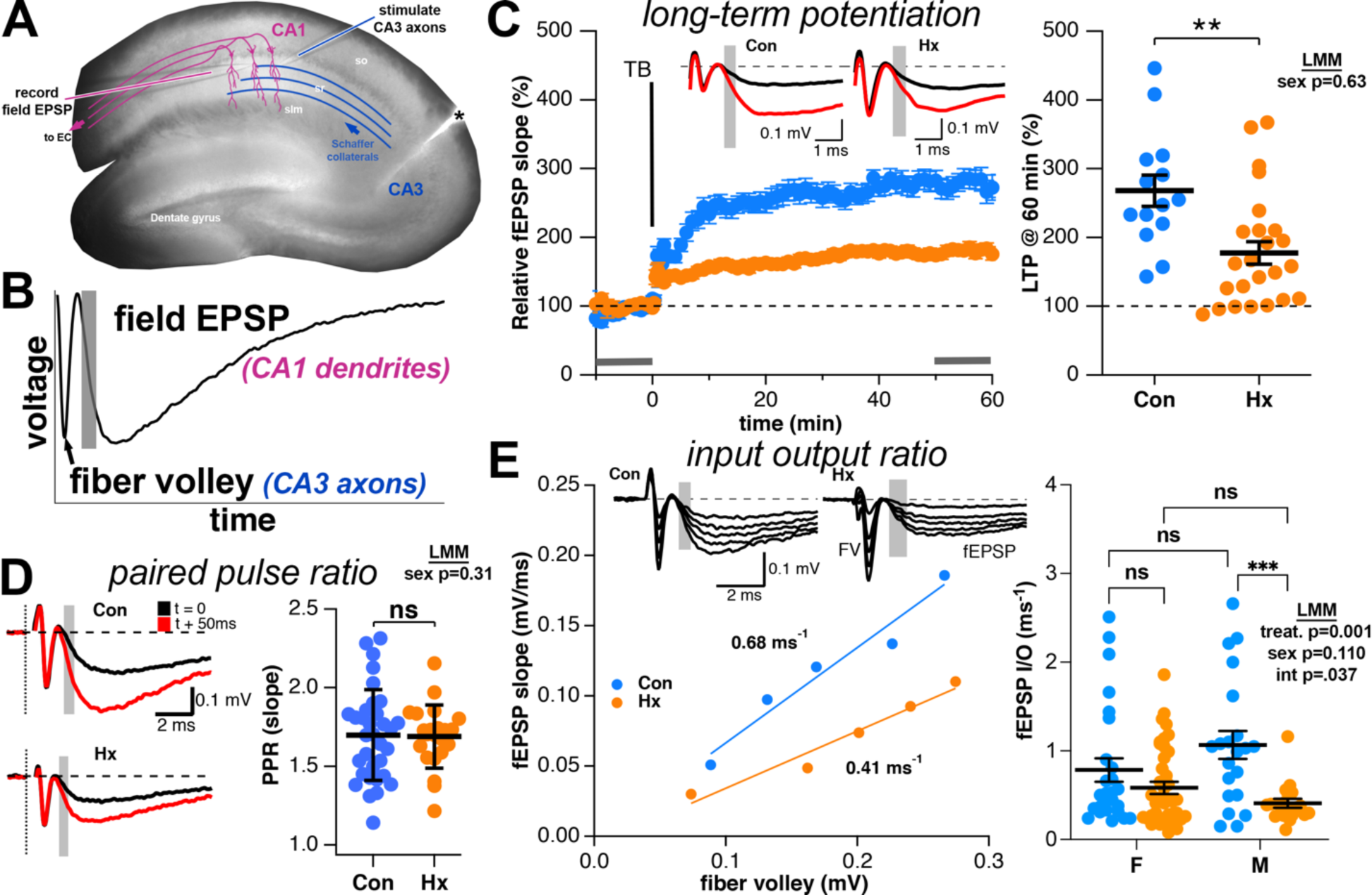
Neonatal Hx reduces adult LTP and synaptic transmission via a postsynaptic mechanism. *A*) Hippocampal slice preparation showing placement of electrodes within the hippocampal CA1 sr and cut made to recurrent CA1-CA3 to block overexcitation (*\**). *B*) Sample field tracing showing the fiber volley (FV, reflecting number of CA3 axons that fired) that precedes an evoked fEPSP (arising from CA1 postsynaptic response). *Shaded area*: initial slope of EPSP reflects predominantly fast acting AMPAR. *C*) Time course of the relative fEPSP slope shows reduced LTP after theta burst (TB) induction. *Inset*: representative tracings before (*black*) and after (*red*) LTP induction. Gray bars highlight region of fEPSP slope measurement. Scatterplot (*right*) showing reduced LTP in Hx (n = 15 slices, 10 mice) compared to control (n = 21 slices, 12 mice, LMM). *D*) Paired pulse ratio, an estimate of presynaptic release probability, is not altered after Hx (n = 22 slices, 4 mice) vs. Con (n = 33 slices, 16 mice, LMM). E) CA3:CA1 input output (I/O) relationship is reduced after neonatal Hx. fEPSP and FV responses to increasing stimulation (*inset*). Representative linear fits of the fEPSP slope:FV relationship in Con vs. Hx. The slope reflects the I/O relation of synaptic transmission. *Right*: Scatter plot of fEPSP I/O from individual experiments analyzed for sex for Con (n = 50 slices, 20 mice) and Hx (n = 63 slices, 18 mice, LMM with Sidak). mean ± SEM **p<0.01, *** p < 0.001.

### Neonatal Hx contributes to altered synaptic strength at CA3-CA1 synapses

To determine whether neonatal Hx disrupts adult CA3-CA1 synaptic strength, fiber volleys (FVs) reflecting presynaptic CA3 axon potentials and postsynaptic fEPSPs were determined (Figs. 4A, B). We first determined if Hx altered presynaptic mechanisms by determining glutamate release probability. Extracellular paired pulse ratio (PPR) measurements were performed by measuring the fEPSP responses to two stimuli 50ms apart, which results in increased glutamate release probability after the second pulse due to residual presynaptic Ca^2+^.^55–57^ The PPR slope was not affected by Hx (Fig. 4D; Con: 1.70 ± 0.05, n = 33 vs Hx: 1.82 ± 0.13, n = 24). Similarly, no differences were found between Hx and Con in other presynaptic factors including the relationship(s) between FV, stimulus intensity and axonal conduction velocity (Supp. Fig. 2). No sex differences or interactions were identified in LTP or PPR responses.

Synaptic strength, as determined by the CA3-CA1 input-output (I/O) relationship, was measured from the relationship between the maximal fEPSP slope and the FV, which reflects postsynaptic AMPAR activation and minimizes postsynaptic voltage-dependent and FV effects.^58,59^ As shown in the figure 4E plot, the individual fEPSP I/O from stimuli between 20 µV to 100 µV were fit with a linear function. This function reflects presynaptic action potential-evoked glutamate release and field postsynaptic membrane response, an estimate of population synaptic strength. Synaptic strength was significantly reduced in animals exposed to neonatal Hx (Fig. 4E scatterplot). In contrast to LTP, synaptic strength displayed a significant interaction between Hx exposure and sex (Fig. 4E). As shown in the figure 4E scatterplot, the effect of Hx was only significant in males after multiple comparison corrections. Thus, the Hx mediated reduction in synaptic strength evaluated by CA3-CA1 I/O appears to be related primarily to sex dependent disturbances in postsynaptic excitatory responses mediated primarily by AMPAR.

### Reduced synaptic density accompanies reduced hippocampal synaptic I/O

Since we did not observe disrupted presynaptic glutamate release probability (Fig. 4D), reduced synaptic strength may be due to either a reduced number of CA3:CA1 synapses or reduced postsynaptic excitatory response. To determine if reduced field synaptic I/O is due to a reduced number of synapses between Schaffer collateral axons and CA1 apical dendrites, we first determined the density of synaptic spines on apical dendrites as an estimate of synaptic density. Using the Golgi staining technique^29^ we determined the density of dendritic spines on the first 25 μm of third-order terminal branches between 100 and 200 μm from the soma (Fig. 5A), the region of maximal arbor complexity and synaptic density within the s.r.^60^ While we identified a significant effect of sex on spine density, no effect of Hx on spine density was identified (Fig. 5B). Reduced I/O may also be due to reduced AMPAR expression. Therefore, we quantified the effect of Hx on excitatory AMPARs and NMDARs by Western blot. Whole hippocampal lysates were generated from young adult animals exposed to Hx and Con littermates and probed for GluAR1, GluAR2, GluN2A, and GluN2B, the primary AMPAR and NMDAR subtypes expressed in mature hippocampal pyramidal neurons.^61^ Unexpectedly, we found no change in AMPAR or NMDAR expression after Hx (Fig 5C). Thus, neither apical spine density measurements nor hippocampal glutamate receptor protein levels were sufficient to explain Hx mediated reduction in I/O.

**Figure 5:**
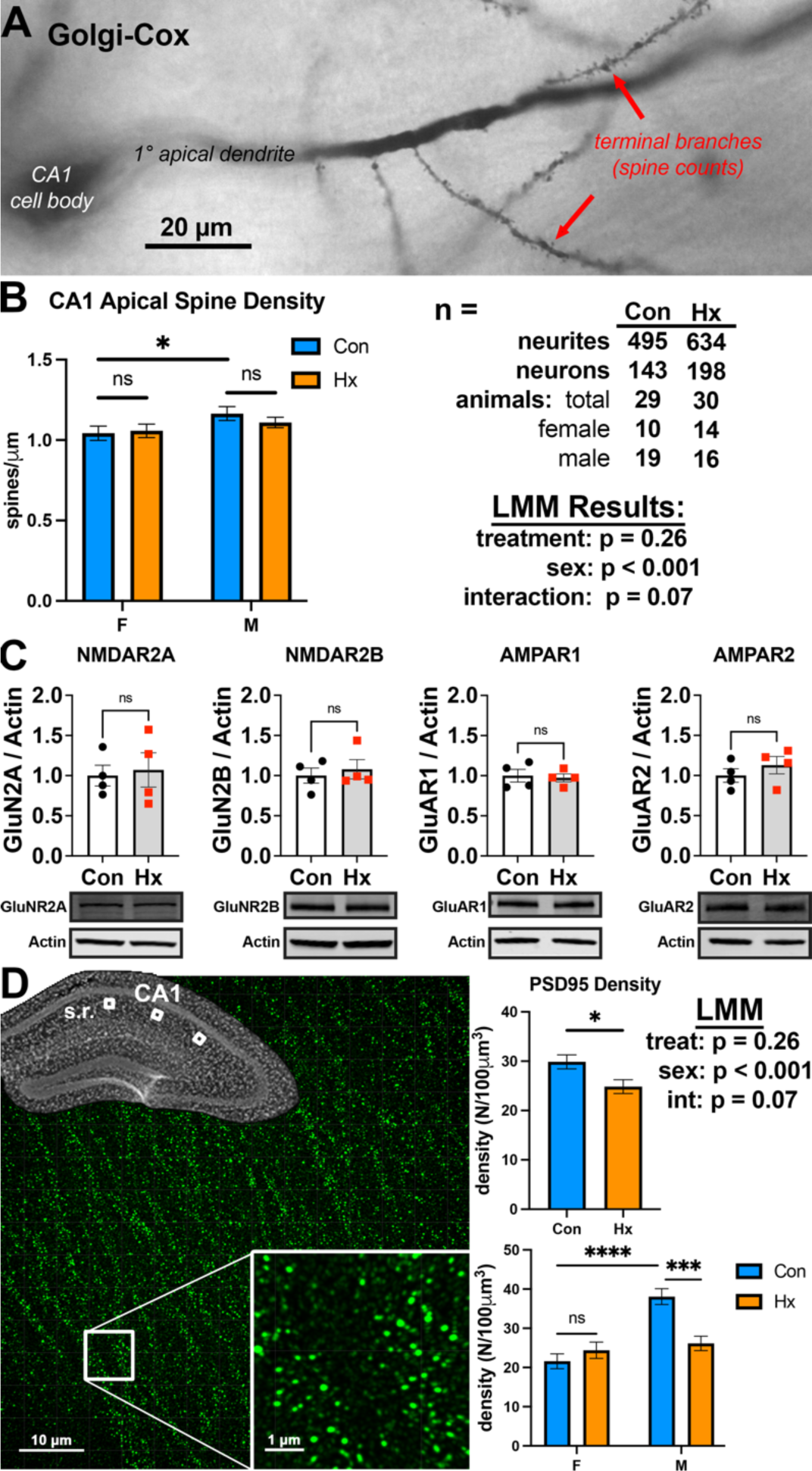
Hx reduces synaptic density in hippocampal CA1 stratum radiatum (sr). *A*) Golgi-Cox filled CA1 neuron with terminal branches showing spine filling (*arrow*). *B*) Apical spine density showed sex difference but no effect of Hx exposure (EMM ± SEM, LMM w/ post hoc Sidak). *C*) Western blots from hippocampal lysates for the primary AMPAR and NMDAR subtypes expressed in adult CA1 neurons were not altered after Hx (n = 4/group, students t-test, mean ± SD). *D*) 3D super-resolution images of immunofluorescent PSD95 puncta were obtained from 3 levels regions of the CA1 sr (*boxes in inset*). Inset at lower right shows detail of individual PSD95 puncta. E) Analysis of PSD95 revealed a loss of synaptic density in Hx (n = 47 ROI, 8 mice, 4 f, 4 m) vs. Con (n = 51 ROI, 8 mice, 4 f, 4 m) mice (*top graph*, EMM ± SEM, LMM) that was most significant in male animals and confirmed baseline sex differences seen with Golgi (*bottom graph*, LMM with Sidak). Regions were pooled after initial modeling showed no differences between regions. * p < 0.05, *** p < 0.001, **** p< 0.0001.

Because spine density as determined by Golgi stain is known to underestimate total spine density due to the challenges of incomplete resolution/filling of small spines and spines oriented in the Z-plane^62^ CA3-CA1 synaptic density was quantified with improved spatial resolution of postsynaptic elements utilizing structured illumination microscopy (SIM, Zeiss Elyra 7) combined with Synaptic Evaluation and QUantification by Imaging Nanostructure (SEQUIN) analysis.^38,39^ This technique has achieved synaptic density quantification equivalent to EM techniques.^39^ To evaluate for regional differences across the CA1,^63^ we analyzed image stacks of PSD95 staining from the CA1 close to subiculum, mid CA1 and CA1 near the CA2/3 boundary from Hx (n = 96 3D image stacks from 8 animals, 4m, 4f) and Con (n = 96 3D image stacks from 8 animals, 4m, 4f) (Fig. 5D). PSD puncta were quantified within each image using Imaris (Bitplane) analytical software. LMM analyses showed no regional differences in synaptic density and data were pooled. In contrast to spine density measures, there was a significant overall reduction in PSD puncta after Hx exposure (Fig. 6D top graph). Furthermore, this was sex-specific, with a greater overall baseline density in males as well as a significant reduction in density due to Hx exposure in male but not female animals (Fig. 6D lower graph). The field I/O relationship (Fig. 4E) had a similar pattern with a Hx-mediated reduction primarily in male animals. This data supports that reduced synaptic density likely contributes to reduced hippocampal CA3-CA1 field I/O after Hx in male mice.

**Figure 6:**
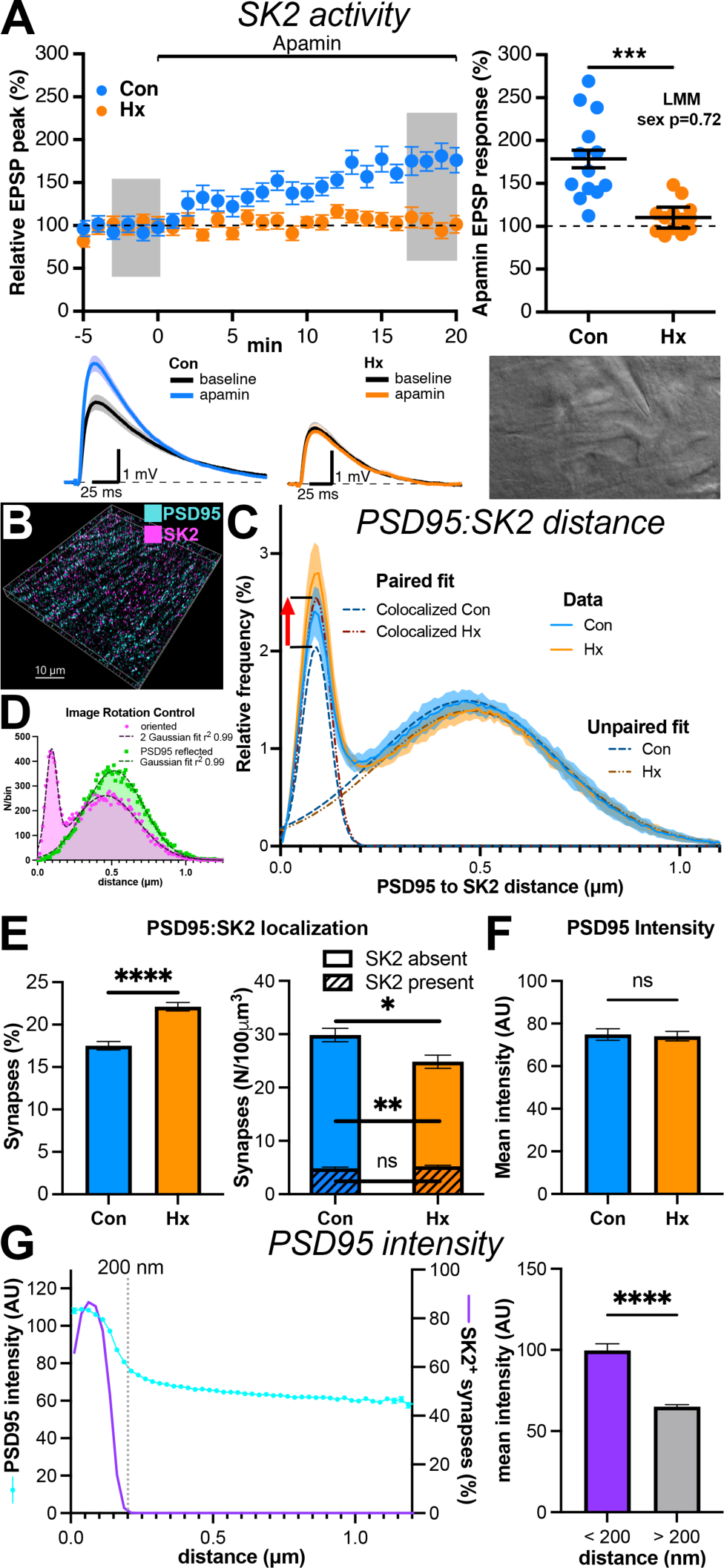
Hx abolishes synaptic SK2 channel activity in adult mice but SK2 channel protein persists at the synapse. *A*) Time course plots of normalized EPSP amplitude of hippocampal CA1 whole-cell recordings (*image*) showing that addition of the SK2 inhibitor apamin (100 nM) increased EPSP in Con but not Hx mice. *Below*: Representative voltage traces for apamin addition in Con and Hx. Scatter plot (*right*) of apamin boosting of EPSP. Con (n = 15 slices, 10 mice, 4 f, 6 m) and Hx (n = 13 slices, 8 mice, 3 f, 5 m). *B*) 3D super-resolution volumes of co-labeled PSD95 and SK2 were obtained from the sr of CA1 hippocampus. C) The relationship between PSD95 and SK2 was defined using nearest neighbor analysis of puncta from image data (mean ± 95%CI) followed by 2 Gaussian fit (*dashed lines shown for each distribution*). The left tall peak corresponds to synaptic pairing of SK2 with PSD95. The red arrow shows the significant increase in the relative frequency of SK2:PSD95 pairs after Hx. The mean and width of the peak remain stable, indicating a stable distance between the PSD and SK2. The wide peak represents random association. *D*) 90° rotation of SK2 in the x-plane caused bimodal PSD95:SK2 data to take on a single Gaussian distribution, confirming the structural relationship of the near peak. *E*) Quantification of PSD95 (synaptic) loci unexpectedly revealed that Hx increased the fraction of synapses containing SK2 (*left*, 1.7×10^6^ pairs, 98 ROI, 16 mice, 8 Con, EMM ± SEM). Data normalized to synaptic density (*right,* students t test mean ± SEM) reveal that synapses without SK2 were lost while SK2 containing synapses were stable after Hx. *F*) Hx does not alter PSD95 intensity (students t test, mean ± SEM). *G*) PSD95 intensity drops when SK2 is between 100 and 200 nm from the PSD center (*blue line*, mean ± SD) as the synaptic SK2 fraction falls (*purple line*, calculated mean). *Right*: Quantification of PSD95:SK2 distance and intensity relationship (students t test, mean ± SEM) shows that SK2 is likely associated with larger PSD regions. No effect of hypoxia on this relationship was found (2 way ANOVA, *not shown*) * p < 0.05, ** p < 0.01, *** p < 0.001, **** p < 0.0001.

### SK2 function is abolished after neonatal Hx

SK2 is a developmentally regulated Ca^2+^ activated K^+^ channel that acts as a negative feedback regulator^21,22^ to limit synaptic depolarization by AMPAR and NMDAR and undergoes activity-dependent trafficking following LTP induction to further increase EPSPs.^54,64–67^ During synaptic transmission, Ca^2+^ influx through the NMDAR activates SK2, which serves to repolarize the membrane potential and reinstate the Mg^2+^ block of the NMDAR thereby modulating neurotransmission and the induction of synaptic plasticity.^22,54^ Altered SK2 function has been implicated in altered hippocampal learning after ischemic injury^68^ and in rodent models of neurodevelopmental disorders.^34,69^ To determine if synaptic SK2 function is altered after Hx, we utilized whole cell patch clamp recordings of evoked EPSPs at CA3-CA1 synapses (Fig. 6A image) before and after bath application of the bee venom apamin (Fig. 6A, diary plot), which specifically blocks SK2. Blocking SK2 with apamin increased evoked EPSPs in Con but not Hx animals (Fig. 6A, scatterplot).^22^ This finding indicates that synaptic SK2 channel function is abolished in young adult mice after neonatal Hx. This loss of function at active synapses may be due to loss of expression, altered localization, or altered regulation.

### Loss of synaptic SK2 function is not mediated by reduced synaptic SK2 expression or localization

SK2 is expressed in multiple neuronal compartments as well as multiple non-neuronal cell types. Therefore, to specifically determine if loss of synaptic SK2 is related to a process that impairs protein expression or synaptic targeting, we utilized SIM microscopy and the SEQUIN analysis to evaluate if Hx alters the nanoscale relationship between the postsynaptic marker PSD95 and SK2. 3D Images were acquired of immunolabeling of both SK2 and PSD95 (Fig. 6B). Imaris software was utilized to identify the 3D location of all labeled puncta within an image. Nearest neighbor analysis was performed as described,^38^ which revealed a bimodal association between PSD95 and SK2 that reflected the population of synapses associated with SK2 (Fig. 6C, narrow paired fit peak) or not (far unpaired pit peak). We found that synaptic (PSD95:SK2) colocalizing puncta were distributed within 250 nm consistent with prior synaptic studies.^39^ We confirmed colocalization by reflecting the SK2 image across the x-axis and performing nearest neighbor analysis.^39^ Altering the orientation abolished the narrow left peak and resulted in a single Gaussian distribution (Fig. 6D), confirming the structural relationship of the particles. Analysis of the fraction and distribution of synapses was performed by fitting the PSD95:SK2 frequency/distance plots with 2 Gaussian distributions and analyzing each distribution. Fit for all data sets were good with R^2^ > 0.98 (e.g., Fig 6C). This analysis unexpectedly revealed that Hx increased the percentage of synapses that co-labeled with SK2 (Fig. 6E % graph). When normalized to the absolute synaptic density defined by the density of PSD95 labeling, however, we observed no change in the density of synapses containing SK2 (Fig. 6E hashed bars), but rather a decrease in those that do not label with SK2 (Fig. 6E open bars). Thus, SK2 appeared to remain in close proximity to postsynaptic elements after Hx.

In addition, the mean and standard deviation of the co-labeled peak reflects the structural relationship between the two proteins. Following induction of LTP, synaptic SK2 are internalized within minutes to inactivate SK2 and potentiate postsynaptic EPSPs.^54,66^ Analysis of the distance between localizing particles was performed to evaluate if SK2 internalization contributes to loss of SK2 activity after Hx. SK2 distance from the PSD center in Con was 89 ± 35 nm (mean ± SD), which agrees with EM-defined PSD95:SK2 relationships,^54,70^ and was not altered after Hx (90 ± 36 nm, Fig. 6C near peak). In order to confirm sensitivity of SEQUIN to SK2 internalization, hippocampal slices were treated with chemical LTP and analysis of PSD SK2 relationship was performed. Although imaging conditions were inferior due to 300 μm sectioning for live hippocampal preparation, the colocalized SK2 peak shifted from 174 to 234 nm after chemical LTP induction (Supp. Fig. 2). Thus, SEQUIN analysis is able to detect channel internalization as previously described,^54,70^ further supporting that no significant SK2 internalization contributes to loss of SK2 synaptic activity.

Intensity based analysis of PSD95 puncta was performed to evaluate for altered postsynaptic structure after Hx. No difference in staining intensity due to Hx was observed (Fig. 6F). Unexpectedly, a strong relationship was found between SK2 distance and intensity (Fig. 6G). Based on data modeling (Fig. 6C), 61% of PSD puncta within 200 nm of an SK2 particle are co-synaptic with SK2 and contain 99.7% of colocalizing SK2 particles (Fig. 6G. line plot). Analysis of this group of SK2-associated synapses showed that PSD95 staining was significantly more intense than synapses much less likely to colocalize with SK2 (Fig. 6G bar graph). Thus, synapses containing SK2 had significantly higher PSD95 staining intensities than those not associated with SK2, which suggested that larger, stronger and more stable synapses^71–73^ are more likely to contain SK2. These data support that SK2 remains expressed within synapses after Hx despite loss of function. In fact, it appears that SK2 associated synapses are relatively preserved after Hx compared to those that are not associated with SK2, despite the loss of SK2 function.

### Loss of synaptic SK2 function is mediated by increased CK2 phosphorylation of synaptic calmodulin

SK2 channels form a complex with calmodulin (CaM), which confers its sensitivity to Ca^2+^. SK2 also interacts with the serine/threonine kinase CK2 and protein phosphatase 2A (PP2A), which regulate CaM Ca^2+^ sensitivity (Fig. 7A, B).^74–76^ CK2 phosphorylation of SK2-bound CaM decreases SK2 opening to Ca^2+^, which reduces SK2 current and has been implicated in neuropathological disease states.^34,77,78^ In order to determine if altered CK2 activity contributes to reduced SK2 function after Hx, hippocampal slices were perfused with the selective CK2 antagonist 4,5,6,7-Tetrabromobenzotriazole (TBB, 10 µM)^34,79–81^ and evaluated for apamin sensitivity. TBB exposure restored apamin sensitivity in cells from Hx exposed animals (Fig. 7C, D). To test if CK2 mediated CaM phosphorylation is altered after Hx in vivo, we directly visualized phosphorylated Thr^79^, Ser^81^ calmodulin (pCaM) in the hippocampi of Hx and Con littermates.^34,81,82^ Because we have seen differential regulation of cellular compartments, we evaluated each layer of the CA1 hippocampus independently. We found that pCaM was increased in the stratum radiatum and stratum lacunosum-moleculare after Hx (Fig. 7E). Thus, Hx-exposed animals increased CK2 phosphorylation of CaM resulting in decreased SK2 open probability and function. Because we saw strong layer variability in the hippocampus we evaluated if a similar response occurred in the cerebral cortex. We observed that pCaM was reduced in somatosensory cortical layers 5 and 6 (Fig. 7F), in contrast to increases in the CA1 hippocampus.

**Figure 7:**
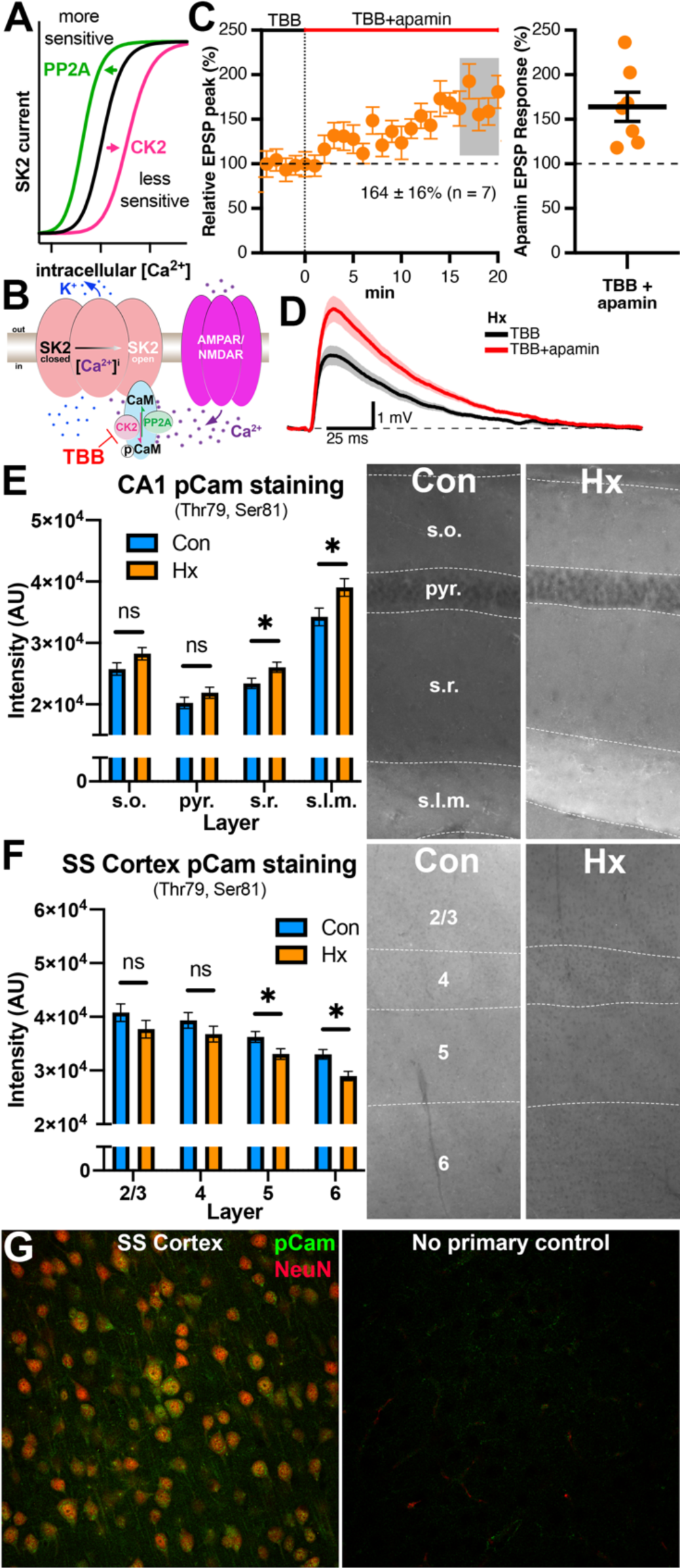
Block of CK2 with TBB restores synaptic SK2 activity. *A*) The kinase CK2 and phosphatase PP2A alter Ca^2+^ dose-response relationships of SK2 via SK2 associated calmodulin. CK2 decreases Ca^2+^ sensitivity, whereas PP2A increases Ca^2+^ sensitivity. *B*) Diagram of the SK2 and associated modulatory proteins including the mechanism of the specific CK2 inhibitor TBB. *C*) Summary time course plots of normalized EPSP amplitude of hippocampal slices from Hx mice bathed in 10 µM CK2 inhibitor TBB. Inset: Representative voltage traces from TBB exposed slices from Hx mice before and after addition of apamin (100 μm). Right: Scatterplot of data from individual experiments (n = 7, mean ± SEM). *D*) Trace of the EPSP response before and after addition apamin in TBB treated slices. *E*) CK2 mediated CaM phosphorylation in the hippocampus (visualized by phosphorylated Thr^79^, Ser^81^ Cam [pCam] antibody, *images*) was layer specific and Hx induced an increase in the sr and slm. *F*) Unexpectedly, CK2 phosphorylation of CaM was reduced in the somatosensory (SS) cortex in a layer specific manner. *G*) Punctate cortical pCam protein (green) was detected in neuron (NeuN, red) cell bodies and associated processes at high power while no primary controls showed no specific staining. * p < 0.05

## Discussion

Although nearly half of preterm survivors display persistent neurobehavioral dysfunction without overt gray matter injury, the underlying mechanisms of neuronal or glial dysfunction remain poorly defined. Despite the fact that preterm survivors are also at risk for cerebral white matter injury that targets oligodendrocyte progenitors, the contribution of dysmyelination also is unclear. To better define the pathogenetic mechanisms that underlie the broad spectrum of behavioral disturbances in these children, we developed a novel model to define maturational responses to cerebral hypoxia in the preterm equivalent P2 mouse. We unexpectedly found that a clinically relevant mild brief hypoxic exposure in the preterm neonate is sufficient to produce hippocampal learning and memory deficits and abnormal maturation of CA1 neurons that persist into adulthood.

It should be emphasized that neonatal P2 mice are much more resistant to hypoxia compared to more mature animals, even under conditions of severe hypoxia.^19,36^ We reported that, when combined with ischemia, P2 mice required a more prolonged exposure to 8% inspired oxygen to generate a combination of gray and white matter injury. Hence, the lack of overt cellular degeneration in our model is consistent with the mild nature of the insult, which was well tolerated without bradycardia, hypoglycemia, severe oxygen desaturations or long-term growth impairment. Thus, our model generates relatively mild hypoxic conditions, which are relevant clinically to sick premature neonates.

Mild hypoxia triggered relatively subtle morphometric and functional disturbances in neuronal maturation without apparent acute or delayed oligodendroglial or neuronal degeneration, dysmyelination or motor deficits. Moreover, hypoxia did not trigger apparent early or delayed responses of astrocytes or microglia, as is typically observed after hypoxic-ischemic white matter injury. It was thus feasible to reproducibly generate hippocampal dysfunction without accompanying white matter injury or neuronal degeneration. Hence, mild hypoxia produced a limited degree of metabolic or oxidative stress during a period of cerebral vulnerability that favors neuronal dysmaturation over glial or neuronal degeneration. Collectively our findings suggest a novel explanation for the broad spectrum of neurobehavioral, cognitive and learning disabilities that paradoxically occur without overt gray matter injury in a high percentage of preterm newborns.^2,4,5,83^

### Hippocampal maturation is persistently disrupted by transient mild hypoxia

Mild *postnatal* hypoxia alone was sufficient to cause persistent CA1 neuronal dysmaturation, which was defined both by structural and functional disturbances in hippocampal maturation as well as learning and memory deficits. Hypoxia was sufficient to disrupt hippocampal CA3-CA1 synaptic strength and LTP and inactivate calcium sensitive SK2 channels within the synapse that are a key regulator of spike timing dependent neuroplasticity, including LTP.^54,65,84^ It is thus noteworthy that a high percentage of human preterm survivors are susceptible to hippocampal-based learning and memory impairment that begins in childhood and coincides with reduced hippocampal growth that persists into adulthood.^6–9^ The sensitivity of the neonatal hippocampus to mild hypoxia was supported by persistent learning and memory deficits in adult mice that were determined using a well characterized fear conditioning model^25^ which distinguishes hippocampal and amygdalar learning. Contextual memory is a measure of hippocampal function^24^ while cued memory controls for deficits in amygdala function.^23^ Hypoxia exposed neonates showed impaired adult learning to context but not cue exposures, supporting selective hippocampal susceptibility.

Contextual memory deficits were accompanied by striking morphometric disturbances in hippocampal CA1 neuronal complexity. Notably, the adult CA1 apical and basal dendritic compartments displayed distinct long-term responses to neonatal hypoxia. Neurite complexity was consistently increased in the basal compartment, whereas the apical compartment appeared unaffected. Moreover, correlation analysis supported that basal and apical complexity appeared to be independently influenced by hypoxia. The specific mechanisms that drive CA1 neurite complexity are unclear,^85,86^ but likely involve both the specific functional roles of each neuronal compartment^87^ as well as interactions with developmental homeostatic mechanisms that regulate overall hippocampal output. For instance, reduced synaptic strength within the CA1 stratum radiatum from the excitatory CA3 input may be balanced by diverse excitatory basal dendritic inputs and result in basal dendritic expansion in the stratum oriens. Hence, increased complexity of basal dendrites may reflect a homeostatic plasticity response to reduced synaptic density in apical dendrites.^88^

Hippocampal field recordings showed persistently reduced synaptic strength at CA3:CA1 synapses after neonatal hypoxia. Synaptic strength defined by the initial slope of the field EPSP is mediated by fast AMPA current and reflects reduced field AMPAR current. This was accompanied by reduced LTP in the same animals, which did not appear to be mediated by presynaptic mechanisms, as no change in the paired pulse ratio was found. Thus, altered CA1 postsynaptic elements appeared to drive hypoxia-mediated dysmaturation. We therefore evaluated postsynaptic drivers of field synaptic strength, including synaptic density, AMPAR, NMDAR and synaptic potassium channels.

### Role of SK2 in neuronal dysmaturation

We observed Hx-mediated reductions in synaptic density, which were accompanied by disturbances in the synaptic potassium channel SK2. SK2 is developmentally regulated, and its expression shifts from low protein levels with predominately somatic localization (P0-P5) to high levels in dendritic and spine plasma membrane (P30).^89^ In mature synapses, SK2 provides negative feedback to AMPA and NMDA mediated neurotransmission, is associated with NMDA and the PSD, and forms a complex with key regulators CaM, CK2 and PP2A. The hypoxia-mediated loss of SK2 activity in young adult animals did not appear to be explained by loss of synaptic SK2 or redistribution within the synapse. Our synaptic microstructure analysis supported that SK2 remained localized to the spine near the PSD. Therefore, we evaluated molecular mediators of SK2 activity including CaM and CK2. SK2 activity was restored in hippocampal slices from hypoxia-exposed animals by application of the specific CK2 inhibitor TBB. CK2 phosphorylates CaM at phosphorylation sites Thr^79^, Ser^81^, which causes altered CaM modulation of SK2 and results in reduced SK2 calcium sensitivity. Thus, our findings support that reduced SK2 calcium sensitivity is mediated through CaM and leads to loss of synaptic SK2 activity. This was further supported by increased Thr^79^, Ser^81^ CaM phosphorylation within the hippocampal stratum radiatum.

Loss of SK2 activity after hypoxia is more likely related to aberrant developmental activation of CK2^74,78^ rather than loss of PP2A activity. PP2A is a nonspecific phosphatase at the same CaM site as CK2, which increases SK2 calcium sensitivity.^90^ However, no CaM specific PP2A drug activators are available to test the potential roles of PP2A. Notably, the robust apamin response in the presence of CK2 blockade also supports significant basal CK2 activity in Hx animals. Interestingly, a similar mechanism was recently described in a mouse model of stress-induced episodic ataxia due to SK2 functional loss in Purkinje cells.^34^

Evaluation of SK2 regulation after hypoxia showed that phosphorylation of Thr^79^, Ser^81^ CaM varies greatly within layers of the CA1 hippocampus. In order to determine if this phosphorylation was hippocampal specific, we evaluated adjacent somatosensory cortex using the same technique and tissue. Although overall levels in the cortex were higher than in hippocampus, surprisingly, hypoxia resulted in reduced Thr^79^, Ser^81^ CaM phosphorylation specifically in the deep cortical layers. Hence, hypoxia-mediated neuronal dysmaturation appears to be regulated in a region-specific manner that likely involves complex interactions within populations of neurons and between brain regions.

Loss of SK2 activity in response to hypoxia may be a secondary homeostatic mechanism related to reduced synaptic transmission and LTP. Reduced SK2 results in a net increase in synaptic strength due to reduced hyperpolarizing adaptation, which may have a paradoxical effect on LTP. Following induction of LTP, synaptic SK2 channels are internalized to further potentiate postsynaptic EPSPs and increase the LTP ceiling, but SK2 function returns within 2 hours.^54,65^ Since SK2 block decreases the frequency threshold for LTP induction,^65^ we expect that reduced synaptic SK2 activity would lead to increased LTP. This may be a homeostatic response to reduced AMPAR mediated synaptic strength to drive LTP. Future targeted mechanistic in vivo studies will be necessary to determine the net effect of CK2 and SK2 modulation on learning and memory.

### Neuronal dysmaturation occurs independently of cell death

Definition of the pathogenesis of learning and memory deficits in survivors of prematurity has been challenging due to the variable overlap of white matter injury with clinical or imaging markers suggestive of gray matter dysfunction. A key finding from our mouse model is that neuronal dysmaturation can occur independently of white matter injury as defined by primary neuronal or glial degeneration or glial reactivity. The lack of overt neural degeneration in our model is thus notable when compared with other neonatal hypoxia models that have employed more severe conditions.^5^ The potential for hypoxia-mediated neuronal dysmaturation to occur in human preterm neonates is consistent with numerous common clinical scenarios where mild or transient hypoxic insults arise from respiratory compromise, which occurs in the absence of apparent ischemia and is not associated with subsequent MRI-defined gray matter injury.^2,4,5,83^

### Technical challenges to detect neuronal dysmaturation after neonatal hypoxia

Since widely employed markers of overt brain injury do not detect evidence of hippocampal dysmaturation, more sensitive approaches were required to detect neuronal maturation disturbances. The application of super-resolution light microscopy for synaptic analysis was key to understanding the role of altered synaptic density and the mechanism of altered SK2 activity after hypoxia exposure. To define the etiology of reduced excitatory synaptic strength, we determined CA1 spine density using the classic Golgi technique, which revealed no differences in spine density. However, further analysis of synaptic density utilizing SIM and the recently developed SEQUIN technique,^39^ showed reduced synaptic density, particularly in males, that mirrored I/O findings from hippocampal field recordings. SEQUIN was also utilized to determine the mechanism of SK2 activity loss in hypoxic survivors. Importantly, we demonstrated that SK2 protein is preserved in synapses close to the PSD, a key finding that led to evaluation of CaM mediated regulation of SK2. At several key points in the study, classic neurophysiology techniques and standard protein-based tissue assays (e.g., Western blot analysis) lacked sufficient sensitivity to determine the specific and relatively subtle structural or mechanistic relationships that were observed in CA1 neuronal dysmaturation. The utilization of this advanced light microscopic synaptic imaging technique allowed the evaluation of key multiprotein synaptic relationships in millions of synapses within the specific cellular layers that corresponded to functional deficits. This led to increased sensitivity and regional specificity, which was not feasible with standard injury-based techniques.

### Sex specificity of neuronal dysmaturation

Sex-dependent influences on the risks for cerebral palsy and neurocognitive outcomes in prematurely born children have been widely noted.^91^ However, it remains unclear to what degree sex may play a role in the development of psychomotor deficits arising from neuronal dysmaturation after perinatal hypoxia exposure. Our findings are consistent with a previously noted male disadvantage for neurological and cognitive outcomes in preterm survivors. We found sex differences in overall animal growth, CA1 dendritic morphology, spine density, and synaptic density in control animals. While both sexes appeared to develop similar hippocampal learning, LTP and SK2 activity deficits, the exposure to hypoxia caused a greater reduction in synaptic strength and density in males than females. The cause for these variable responses remains unclear, but underscores the strength of our model to replicate influences of sex on human neurodevelopmental vulnerability to preterm birth.

### Pitfalls and future directions

Despite extensive evaluation of the roles of early or late oligodendrocyte injury, glial reactivity and primary neuronal injury, the triggers for persistent neuronal dysmaturation remain unclear. Further studies to evaluate single cell types for genetic or epigenetic modifications, which occur in response to hypoxia may yield additional insights into key mechanisms of initiation and maintenance of dysmaturation. Our study also focused on excitatory signaling changes in the hippocampus. Further studies of the homeostatic relationships between different neuronal compartments and potential compensatory changes in excitatory/inhibitory balance may yield other novel mechanisms that could alter neuronal dysmaturation and potentially restore hippocampal learning and memory function. In addition, we expect that brain regions other than the hippocampus may be vulnerable to neuronal dysmaturation.

## Supporting information

Antibody list

## Acknowledgments

We thank the OHSU Advanced Light Microscopy Core, especially Drs. Stefanie Kaech Petrie and Brian Jenkins for invaluable assistance with structured illumination microscopy. Supported by NS116674 and HL163517 to SAB and CNCDPK12 to AR.

