## Supplemental Table 1 for "Mild neonatal hypoxia disrupts adult hippocampal learning and memory and is associated with CK2-mediated dysregulation of synaptic calcium-activated potassium channel KCNN2"

| **Field property** | | | | | | | | | | | | | |
| --- | --- | --- | --- | --- | --- | --- | --- | --- | --- | --- | --- | --- | --- |
| **Variable** |  | **Significance (p)** | | | **Condition** | |  | | | **Mean^‡^** | |  | **SE** |
| FV vs. stim intensity  (μV/μA) | | | 0.769 |  | Con  Hx | |  | 4.68  4.67 | | | | | 0.66  0.56 |
| Conduction Velocity  (mm/s) | | | 0.189 |  | Con  Hx | |  | 162.2  154.0 | | | | | 4.8  3.9 |

**Supplementary Table 1:** No change in presynaptic factors including fiber volley (FV):stimulus intensity or conduction velocity were found after Hx. FV:Stimulus intensity relationship was plotted from 20-100 μA and fitted with a linear relationship, the slope of which describes CA3 axon recruitment and excitability. Conduction velocity was calculated from the distance between stimulating and recording electrodes and the time between stimulus and FV initiation. (Con n = 50 slices, 22 animals, Hx n = 63 slices, 19 animals) Linear mixed models (treatment, sex, interaction) showed no significant sex effect or interaction; ‡: estimated marginal means are shown
