## Supplemental Figures for "Mild neonatal hypoxia disrupts adult hippocampal learning and memory and is associated with CK2-mediated dysregulation of synaptic calcium-activated potassium channel KCNN2"

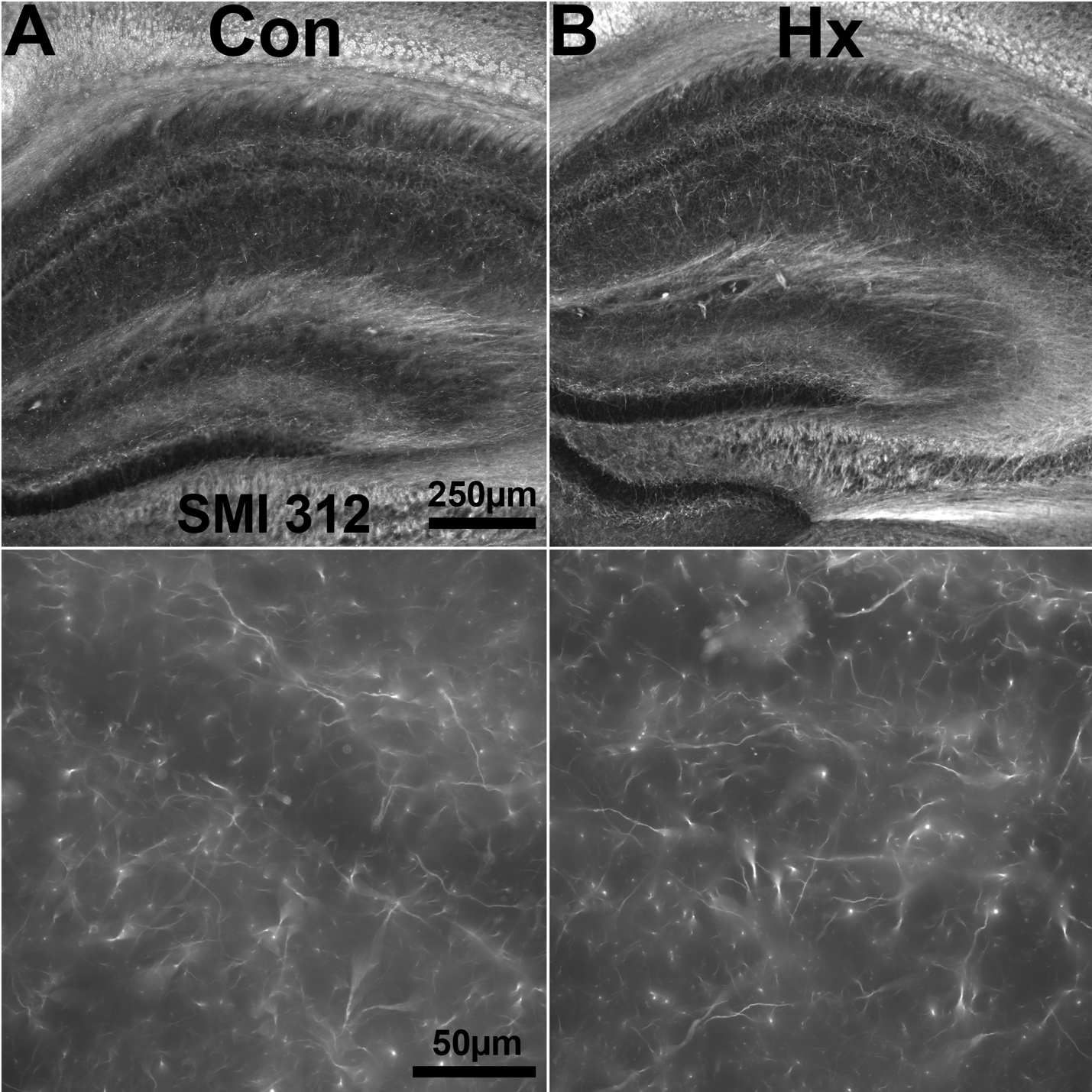

**Supplemental Figure 1:** Axon integrity appeared normal after neonatal Hx exposure. Representative neurofilament staining with SMI 312 in the hippocampus of control (A) and littermate Hx exposed (B) mice at P50. No evidence of axonal injury, including no focal swellings, altered staining intensity, or fragmentation was seen within the hippocampus, WM or corona radiata in Con or Hx animals.

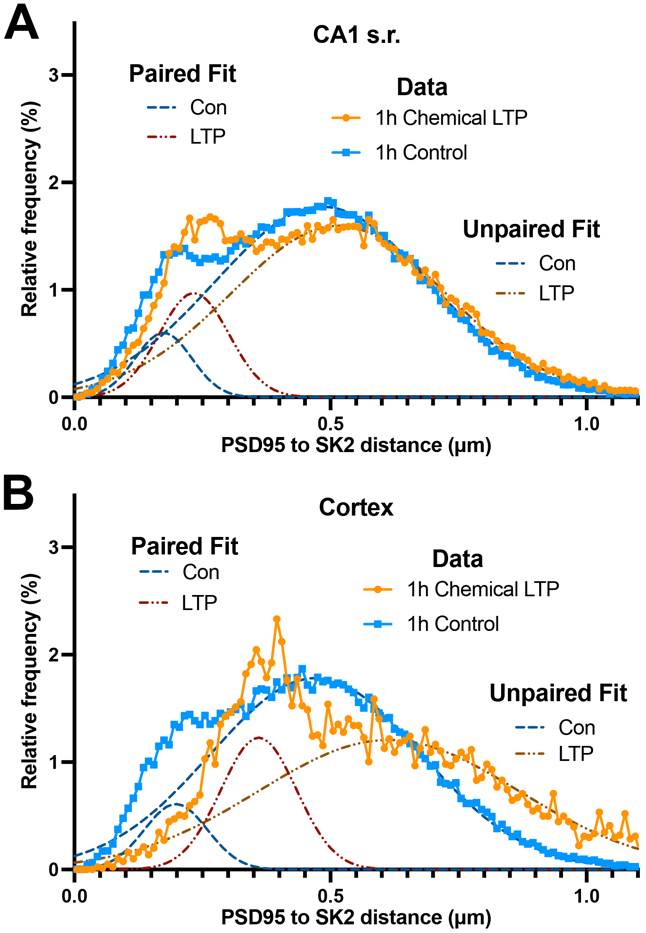

**Supplemental Figure 2:** SIM imaging and SEQUIN processing detected chemical LTP-induced SK2 to shift away from the PSD. 300 μm live hippocampal slices were prepared and underwent chemical LTP induction to evaluate SIM imaging and SEQUIN processing sensitivity. Although signal-to-noise ratios were inferior to 50 μm sections, SEQUIN imaging detected the chemical LTP-induced shift of SK2 out of the PSD in the CA1 hippocampus (A) and cortex (B), as previously demonstrated by electron microscopy.^1,2^
